# Nucleus Basalis of Meynert as a DBS target

**DOI:** 10.1101/2025.06.11.659014

**Authors:** E Levichkina, TR Vidyasagar, C French

## Abstract

Several clinical attempts to use deep brain stimulation applied to nucleus basalis of Meynert to reduce cognitive decline in different types of dementia report inconclusive effects. At the same time, experiments in rodents have largely demonstrated cognitive improvement. The hypothesised basis for this difference was the application of different patterns of stimulation – tonic continuous over multiple sleep-wake cycles in human clinical trials vs intermittent bursts over shorter intervals in animal studies. However, since no systematic testing of the effects of different stimulation patterns applied in different states of vigilance was conducted, it remains unclear what specific attributes of cortical responses are associated with these patterns, and the effect of vigilance state on pattern response.

We hypothesised that communication between nucleus basalis and its cortical targets involves frequencies in both low and high ranges, making burst-type intermittent patterns potentially more effective. Therefore we studied preferred frequencies of neuronal synchronization within nucleus basalis as well as between nucleus basalis and its cortical targets in B6 mouse strain (N=9, age 4-6 month) by recording neuronal spiking and local field potentials (LFP) from 2 sites of the nucleus basalis and 4 cortical sites simultaneously across multiple sleep-wake cycles. Spike-field coherence (SFC) analysis revealed natural synchronization tendencies of nucleus basalis – all animals demonstrated significant SFC at delta-theta range and gamma coherence peaking between 57 and 88 Hz (SFC > 95% Jackknife confidence interval at frequency range > 2*bandwidth). SFC between cortical cells and local field potentials of nucleus basalis, which reflects cortical feedback, peaked at delta-theta frequencies.

Based on these results we constructed “nested” patterns of stimulation that included both delta-theta and individually assessed gamma peak for each animal (N=8). The effects of nested stimulation were compared to the ones of random and tonic (20Hz) stimulation patterns. These patterns of stimulation were applied across multiple sleep-wake cycles to freely behaving animals, and spectral power density in response to stimulation compared between patterns for different states of vigilance.

We report significant reduction of delta power during slow wave sleep and theta power in REM for all three patterns. Slow wave suppression was also present during wakefulness. This indicates all patterns may improve levels of alertness but highlights strong likelihood of sleep disturbances in response to stimulation during sleep. Thus, deep brain stimulation of nucleus basalis may be counterproductive during sleep.

The nested pattern of stimulation alone demonstrated preservation or even enhancement of gamma activity in the cortex while suppressing low frequencies. Broad-band suppression of cortical activity was shown in response to tonic stimulation, and to some extent to the random pattern as well, suggesting that tonic stimulation may be unsuitable for deep brain stimulation of the nucleus basalis of Meynert.

## Introduction

The impressive success of Deep Brain Stimulation (DBS) in alleviating motor symptoms of Parkinson’s Disease (PD) raised the question whether the same technique can be helpful in treating various forms of dementia (Nazmuddin et al., 2021; Picton et al., 2024). Thus, identifying an appropriate DBS target to potentially reduce cognitive decline became of great interest. On the other hand, stimulation of subthalamic area or globus pallidus interna – common targets of DBS in PD - are known to sometimes worsen symptoms of dementia in PD patients (Morrison et al., 2004; Hariz et al., 2008; Rothlind et al., 2015).

Two main targets chosen by clinical trials working on Alzheimer’s disease (AD), Lewy body dementia (DLB) and Parkinson’s disease dementia (PDD) were the fornix and nucleus basalis of Meynert (NBM). Stimulation of the fornix, an important part of memory-related circuits and the main pathway between the hippocampus and subcortical structures such as the anterior thalamic nucleus, was targeted to test effects on memory (Liu et al., 2020; Ríos et al., 2022). The NBM was targeted to possibly enhance cholinergic input to the cortex, hippocampus and amygdala, and due to its known deterioration in AD and other dementias (Gratwicke et al., 2013, reviewed by Nazmuddin et al., 2021). Considering that NBM is the main source of cortical acetylcholine (Ach) (Mesulam et al. 1983; 1988; Liu et al., 2015), the diffuse distribution of its outputs (Sarter et al., 2009) and its ability to influence cortical plasticity via astrocytes (Takata et al., 2011), NBM is a target to provide widespread cholinergic activity and potential modulation of cognitive function and level of arousal, both of which are impaired in dementia.

However, results of DBS clinical trials that targeted NBM showed at best moderate and sometimes inconclusive effects (Nazmuddin et al., 2021). Gratwicke et al. (2018) conducted a small randomised trial in PDD patients and found no significant improvement of cognitive function, with the only positive outcome being decrease in hallucinations, while Freund et al. (2009) demonstrated a marked improvement in attention, memory and alertness in their PDD patient, possibly related to their dual-target approach, where subthalamic nucleus was stimulated to achieve normalization of motor function, while NBM was stimulated to help with cognition. In a study involving AD patients tonic stimulation was able to reduce the speed of the disease progression but only in 2 out of 6 patients (Kuhn et al., 2015). Xu et al. (2024) showed short-term cognitive improvements but no long-term disease modification in a 12-month follow up. These disappointing results are in a striking contrast with a number of animal studies of NBM DBS, which reported improvement of many cognitive parameters (reviewed by Nazmuddin et al., 2021; Kumbhare et al., 2018). We therefore sought to investigate differences in the stimulation parameters that were used in the animal and human studies, in order to optimise the approach to DBS.

One of the key differences is the mode of stimulation and the other is its duration. Tonic prolonged stimulation of 10-50Hz dominated in human clinical trials. On the other hand, since the time of stimulation was limited in the animal experiments where tethering of the animal to the stimulation equipment is necessary, NBM DBS in animal models relied on intermitted and/or burst pattern of stimulation applied for a limited time (Lee et al., 2016; Koulousakis et al., 2020; Kumbhare, 2024; Nazmuddin et al., 2021). That the animal studies were successful in achieving cognitive improvements may have been due to this difference in stimulation pattern and the fact that the stimulations were done during wakefulness.

Transient bursts of activity are more representative of cortical dynamics than sustained activity in many contexts (Buzsáki and Wang, 2012; van Ede et al., 2018; Akam et al., 2014; Vidyasagar & Levichkina, 2019), including memory (Lundqvist et al., 2022) and attention (Anderson et al., 2011). A recent study in rats with saporing-induced dementia demonstrated the superior effect of burst-type stimulation comparing to tonic stimulation in improving results of novel object recognition task and auditory learning (Kumbhare, 2024). Non-Ach cells of the basal forebrain which are linked to motivational salience, have been shown to demonstrate bursting behaviour as well (Lin et al., 2008), while cholinergic cells demonstrated bursts of gamma activity modulated by theta rhythm and corresponding to cortical activation (Lee et al., 2005). In addition, tonic stimulation was demonstrated to decrease performance in a working memory task in a primate model of NBM DBS, while intermittent stimulation resulted in improvement (Liu et al., 2017). At the same time, studies conducted in PD patients showed NBM cells firing at 13 ± 10 Hz during wakefulness, similar to the 10-50Hz used in tonic stimulation (Lee et al, 2020). Rodent measurements of activity in various NBM cell groups showed that Ach cells had slow firing rates (in delta-theta range) while PV GABA-ergic and glutamatergic cells had higher firing rates (beta to low gamma range) (Xu et al., 2015). However, mean firing frequency is not always representative of the synchronization frequency of neurons in the neuronal ensembles they form, and which are more efficient in transmitting information to the target area (Gray et al., 1989; Yu et al., 2004; Fries, 2005; Saalmann et al., 2007; Bastos et al., 2015).

The efficiency of NBM stimulation pattern in changing cortical activity could be linked to natural synchronization frequencies of its cells. Kumbhare et al. (2018) argued that “delta-gamma” or “theta-gamma” burst patterns of stimulation can potentially be more beneficial as they are more likely to engage cortical circuits via synchronisation. We therefore hypothesised that the application of a stimulation pattern based on the natural synchronization peaks of NBM should evoke more prominent wakefulness-related cortical activity in comparison to other patterns.

The other possible issue with the effectiveness of the stimulation in human clinical trials is the prolonged application, for as long as a year through multiple sleep-wake cycles. Stimulation of the main source of cholinergic support to widespread areas of the brain during sleep is likely to erode sleep quality (Irmak and de Lecea, 2014; Nazmuddin et al., 2021; Kumbhare, 2024). AD, PD and DLB all have sleep deterioration as an integral part of the respective clinical pictures. AD patients suffer from general deterioration of both NREM and REM sleep, including decrease of the overall lengths, fragmentation, increases of sleep latency and of the number of awakenings during the night, with the degree of deterioration linked to the severity of cognitive decline (Liguori et al., 2014; Mander et al., 2015; Kuang et al., 2021; Zhang et al., 2022). PD patients often have insomnia often accompanied by other sleep disturbances such as sleep-related breathing disorders, restless legs syndrome and rapid eye movement (REM) sleep behaviour disorder (Menza, 2010; Minakawa, 2022). Lewy body pathology is so tightly linked to REM-sleep behavioural disorder that it is considered to be one of the diagnostic criteria for the disease (Boeve et al., 1998; McKeith et al., 2017). At the same time, excessive daytime somnolence (EDS) is also an important part of pathology in these diseases (Kasanuki et al., 2018; Elder et al., 2022; Carvalho et al., 2018; Feng et al., 2021). It is especially severe in DLB, affecting as many as 90% of the patients (Elder et al., 2022). Therefore, an effective DBS mode of stimulation is likely to be the one reducing excessive sleepiness when the patient needs to be awake when brain activity has less delta activity and more pronounced gamma activity. The very same mode of stimulation is likely to disrupt slow wave sleep (SWS), potentially “cancelling” any positive effects of DBS-related memory improvement during wakefulness due to disturbance of memory consolidation processes depending on sleep (Klinzing et al., 2019; Chang et al., 2025). Furthertmore, it may contribute to cognitive decline due to the severity of sleep disorganization in AD (Liguori et al., 2014; Mander et al., 2015; Kuang et al., 2021; Zhang et al., 2022). It is currently unknown whether NBM stimulation can also be detrimental for REM sleep, which is already severely affected in DLB. We decided to characterise cortical responses to NBM DBS across sleep-wake cycles to establish what kind of alterations of cortical rhythmic activity are caused by different patterns of stimulation in different states of vigilance.

Thus, we aimed to study the effects of various DBS patterns applied at different states of vigilance on the cortical targets of NBM. Understanding the basic nature of these responses in healthy animals would provide a benchmark for the effects of stimulation in normalising cortical activity in disease by DBS. Therefore, we implanted electrodes into NBM and several cortical sites of healthy adult mice and designed two experimental paradigms.

The first one involved simultaneous recording of neuronal activity (single cells and local field potentials – LFP) from NBM and the cortex to determine natural cell-LFP synchronization peaks within NBM, which is necessary to observe NBM output frequencies, as well as between cortical cells and LFP of the NBM, which is needed to characterise cortical feedback frequencies that might affect NBM and reinforce synchronization at certain frequencies. We used spike-field coherence (SFC) as a measure of synchronization.

The second experiment utilised results from the first one to design a DBS pattern matching the natural synchronization peaks for each animal. This pattern was used for NBM stimulation across multiple sleep-wake cycles along with two other patterns: a control random stimulation and a 20Hz tonic stimulation to match the one used in human clinical trials. This allowed us to analyse cortical responses in the frequency bands commonly used to characterise SWS (delta), REM (theta) and wakefulness (gamma) frequencies to describe responses evoked by different patterns of stimulation.

Deterioration of delta waves is a hallmark of SWS disturbance (Long et al., 2021; Verma et al., 2024), while theta waves characterise REM and are essential for hippocampal function (Buzsáki, 2002). Correspondingly, disruption of delta activity in SWS and theta-activity in REM would indicate deterioration of these sleep stages, which is undesirable when a person needs to sleep. At the same time, reduction of delta activity during wakefulness would be a positive sign of the capability of stimulation to reduce daytime sleepiness. However, reduction of slow wave activity needs to be accompanied by an increase of the higher frequency components of cortical activity. Beta and low gamma activities are especially of interest due to their strong link to active conscious behaviour (Bastos et al., 2015; Buschman & Miller, 2007; Fries et al., 1997) and attention (Saalmann et al., 2007; Esghaei et al, 2022). Moreover, low gamma activity deteriorates in AD patients while high gamma changes are less known (Casula et al., 2022; Arroyo-García et al., 2021; Murty et al., 2021). PDD and DBL are characterised more by general lowering of EEG power (Bonanni et al., 2008; van der Zande et al., 2018), with some evidence for low gamma deterioration as well (Güntekin et al., 2023). It was therefore important to find out the kinds of gamma activity changes that occur as a result of NBM DBS. To determine whether DBS affects other physiologically relevant oscillations we included alpha and beta bands in the analysis as well.

## Methods

### Animals

9 male C57BL/6J healthy mice aged 4-6 month were used in this study. All of them were implanted with 6 stimulation/recording electrodes; however, due to deterioration of one of NBM electrodes in one of the animals, stimulation experiment was conducted only in 8 animals, while SFC measures were derived from all 9. After surgery animals were individually housed with access to food and water ad libitum, at a constant temperature (20–22°C) and ventilation, with an automatic 12-hour light-dark cycle.

### Surgery and electrodes implantation

The mouse was anesthetized box with 5% isoflurane in O2. After the animal was unresponsive, it was injected with Meloxicam (5 mg/kg, subcutaneously) and mounted in the stereotaxic apparatus (WPI, Manual Mouse Instruments), with its nose in the nose-cone of the stereotaxic apparatus and the head fixed with the ear bars. A heating pad was used during the surgery to maintain the body temperature. The mouse was deeply anesthetized with 1–2% isoflurane in oxygen during the surgery. Dorsal surfaces of its head and neck area were shaved and then cleaned with 80% ethanol and further disinfected using Iodine and ViralFX™ Pink Solution (Chlorhexidine 0.5% in Alcohol 70%) solutions. An eye ointment (retinol palmitate, VitA-POS®) was applied to the eyes for protection.

For intracranial recordings, we used varnished tungsten electrodes (FHC) with impedances between 0.5 and 1 mΩ. The mouse equivalent of NBM includes the horizontal limb of the diagonal band (HDB), substantia innominata (SI) and Magnocellular preoptic nucleus (MCPO) (Fug 1A). A pair of NBM electrodes with tip separation of 1.1 mm was vertically introduced into the right nucleus basalis at LAT+1.45 mm, AP 0.00 mm, at a depth of 5.61 mm from the dura for the deeper ones and 4.9 for the more superficial ones. The coordinates were chosen using Allen Mouse Brain Atlas (Dong, 2008). The electrodes were introduced simultaneously and positioned in a way, so that the posterior electrode was directed more laterally and more superficially than the anterior one to stimulate a substantial portion of NBM when the current was applied between these two electrodes.

Four cortical electrodes were also implanted into the right hemisphere: Frontal pair of electrodes (ML + 2.76 mm; AP + 1.83 mm) - lower electrode targeted primary motor cortex (M1), the other targeted Anterior insula (Ai); Parietal pair of electrodes (ML + 4.4 mm; AP -1.57 mm) - lower electrode targeted supplementary somatosensory cortex (SS), the other targeted Visceral insula (Vi).

Thus, 3 craniotomies were made for these 3 pairs of electrodes. Since NBM is expected to supply the entire cortex (Fibiger, 1982; Mesulam et al., 1983; Chaves-Coira et al., 2018), potentially any cortical cite can be used to study evoked responses to NBM stimulation. Our choice was guided by the geometry of the mouse brain that restricts accessibility for implantation of the areas placed too closely, as well as the evidence in the literature for the involvement of these areas in dementia (Arnold et al., 1991; Suvà et al., 1999; Roquet et al., 2017; Orta-Salazar et al., 2019; Kitamura et al., 2020; Ferreri et al., 2003; Zhang et al., 2023).

EEG electrode (1.2 M stainless steel screw) was implanted in the skull, overlaying the hippocampus, to be sensitive to the theta oscillation during REM sleep (ML -2.01 mm, AP -2.48 mm). The reference/ground screw was positioned posteriorly over the cerebellum area. These screw electrodes were placed epidurally over the left hemisphere. Two electromyographic (EMG) hook electrodes (MicroProbes, Stainless steel, 50-micron diameter) were introduced into the neck muscle through a 27G needle.

All electrodes were connected to a prefabricated female connector after electrode implantation. The area was sealed with CA glue (BSI INSTA-CURE^+™^) and secured with dental acrylic cement (PYRAX^®^, rapid cure acrylic) to ensure the structural integrity of the implant. The overall structure enclosing the electrodes and containing the connector will be referred to as the headpiece. The animal received Meloxicam subcutaneously (5 mg/kg BW) in the postoperative period every 12 h for up to 48 h for pain relief and was provided with soft food (peanut butter, fruit mash) and sunflower seeds during the recovery period. A recovery period of at least 1 week was used before the onset of the recording experiments. After recovery from surgery, the animals were gently handled for 3 days and brought into the recording room to accustom them to the recording situation. After that, the headpiece connector was attached to the Intan lightweight cable counterbalanced above the animal’s cage in a way allowing the animal to move freely (Figure 1B). After the animal showed the ability to fall asleep while the headpiece was connected (which took 1-3 days of training), the experiments involving recording of brain activity could commence.

**Figure 1.**
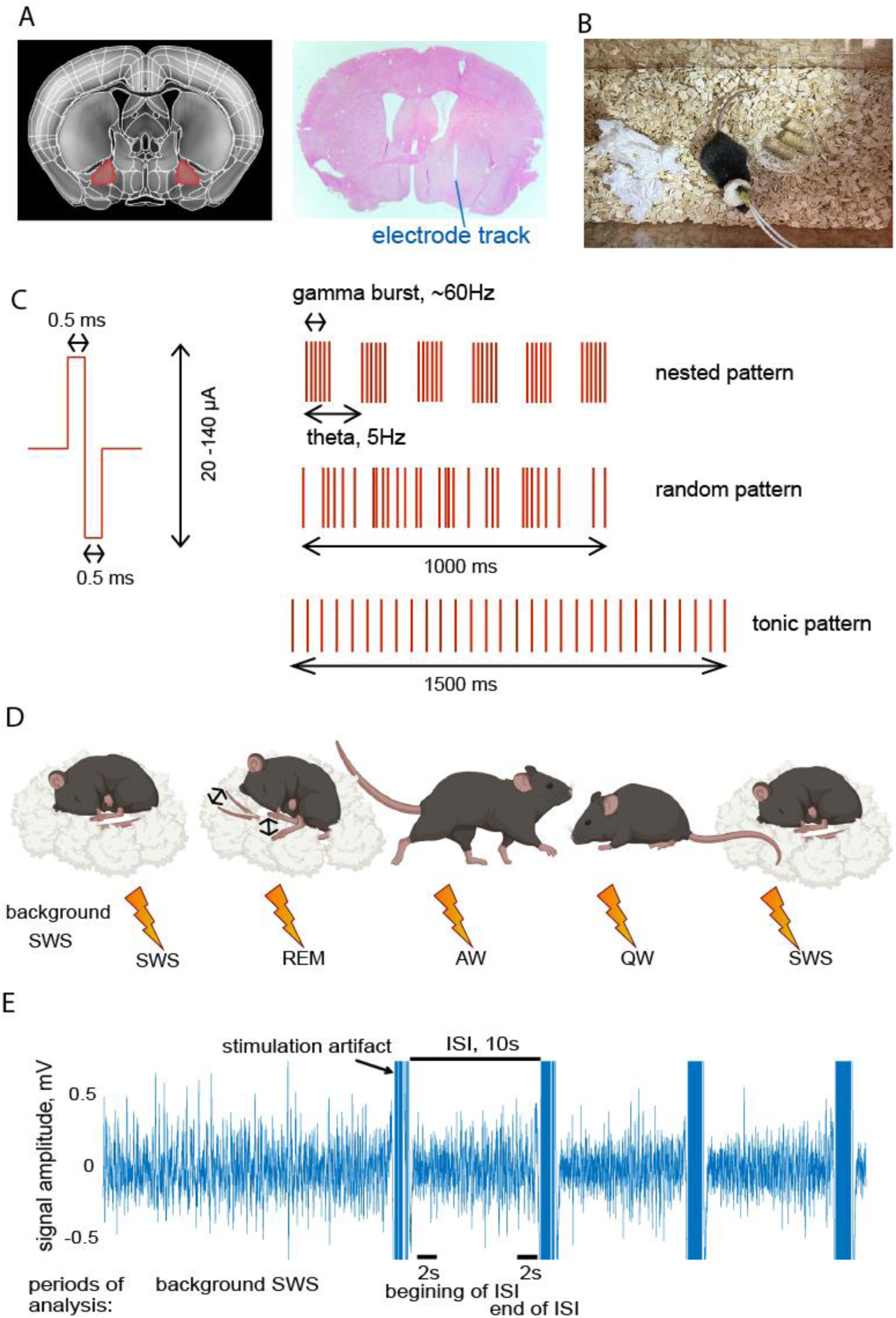
**Methods** A. NBM area according to the Allen Mouse Brain volumetric atlas (Dong, 2008) on the left, with the target area highlighted by red, and a histological section with the NBM electrode track from one of the animals on the right. B. Video monitoring of the animal during one of the recording sessions. C. Biphasic pulse and patterns of stimulation used in Experiment 2 – nested, random and tonic. D. Schematic depiction on a single recording session of Experiment 2 that started from recording of the slow wave sleep (SWS) background, after which stimulations were done across all states of vigilance (starting in SWS as well): Rapid Eye Movement sleep (REM), Active wakefulness (AW), Quiet Wakefulness (QW) (see detailed description below). Created using Biorender.com. E. Raw data example from a single channel: background SWS followed by stimulation. A decrease of slow wave power can be seen after every individual stimulation.

### Electrodes positions confirmation

After data collection (see below), the animals were euthanized with a 2% lethobarb injection, immediately followed by transcardial perfusion with phosphate-buffered saline (20 mL PBS/ea), then by 4% paraformaldehyde (20 mL PFA/ea). Brain tissue was histologically examined to confirm positioning of the NBM electrodes using either Hematoxylin and Eosin or Crecyl violet staining.

We report here the results obtained from the animals having stimulating electrodes positioned in the NBM area. Figure 1A demonstrates the NBM area according to the Allen Mouse Brain volumetric atlas (2013) on the left and a histological section with the NBM electrode track from one of the animals on the right.

### Data acquisition

Recordings were conducted during daytime across several sleep-wake cycles (Fig.1, panels B and D). Individual recording sessions lasted up to 2h, and sessions were conducted with at least one day interval between them. All recordings were done while the animal stayed in its own cage to reduce any stress. The cage was placed inside a Faraday cage made of copper mesh and grounded to the data acquisition system to reduce noise. Intan RHS acquisition system was used for both recording and stimulation. The hardware amplifier bandwidth was set at 0.5-5000 Hz, and all signals were sampled at 20000 Hz per channel. An impedance test was done for each channel before the onset of recordings to make sure the normal functioning of each electrode.

### Categorization of the states of vigilance

To avoid disturbing the animal’s normal sleep-wake cycles, we used videomonitoring combined with visualised brain activity and EMG during the recordings for preliminary classification of the animal’s activity as putative wakefulness, SWS or REM and recorded observations. Later, we manually categorised recordings off-line and marked four states of vigilance: active wakefulness (AW), quiet wakefulness (QW), slow wave sleep (SWS), and rapid eye movement (REM) sleep, according to the EEG oscillation patterns and motion activity observed via EMG signals as well as the protocol of the animal’s behaviour during the session. Sections of recording containing motion artifacts were not included in the analysis. The reason behind subdividing the state of wakefulness into AW and QW was to prevent averaging neuronal spike rates between seemingly different states as neurons can give a short burst of spiking in one state, e.g., during specific activity, that does not persist for a long time. To distinguish the AW and QW, we defined a period of less than 5 sec from the offset of a vigorous movement as AW, and a period further than 5 sec from such a movement as QW as long as no noticeable delta waves were present in the EEG and the animal was not behaviourally asleep.

Categorisation was done by placing markers corresponding to a particular state of vigilance at intervals of >3 sec from each other for further analysis. Transition periods and intervals containing motion artifacts were not marked and therefore not included in further analysis. Recording duration was deemed sufficient if all 4 states of vigilance yielded the number of 3 sec intervals needed for statistical analysis. Commonly, the bottleneck for collecting enough data was an AW state without motion artifacts during each of the 3 sec intervals. Achieving >30 AW such intervals also ensured collection of sufficient amount of data belonging to other states. If that was not possible within a daily session, several sessions were conducted and combined.

### Experiment 1. Determining natural frequency peaks of cell-LFP synchronization

At least 2 recording sessions without stimulation containing all four states of vigilance were done and visually examined to establish sections of recordings corresponding to each of the 4 states (Fig. 1, panel D). Data from these recordings were used for SFC analysis.

Raw signal recorded from each of the implanted electrodes was band-pass filtered (Butterworth, 2-order, bi-directional) at 300-3000 Hz for further spike sorting and at 0.5-250 Hz (and resampled at 1000 Hz) for LFP analysis.

As many neurons can be present near the tip of the electrode, individual neuronal signals are commonly spike-sorted off-line using algorithms that estimate spike amplitude, polarity, and waveform shape to determine the similarity between spikes and decide whether they belong to the same or different cells. To this end, we used the inbuilt algorithm of Spike 2 software (Cambridge Electronic Design Limited; for further details and an illustration of the process, see e.g. Levichkina et al., 2021). These algorithms usually cannot separate data from all of the many cells surrounding the electrode due to algorithm saturation (Pedreira et al. 2012) and the number of correctly defined neurons can potentially range between 1 and 9, in our hands. However, the yield of well-separated spikes can be optimised or even increased if a particular recording site demonstrates clusters of neuronal activity of distinct amplitudes. In such a case the upper and lower amplitude thresholds can be applied to separate these different clusters and the spike sorting run separately for each of the clusters. This situation is especially common in the brain areas containing several types of neurons having very different sizes, such as NBM. As Spike 2 offline spike sorting provides the option fo spike sorting within such different thresholds, we used this approach for the recording channels that demonstrated such clusters, as exemplified by Figure 2; therefore yield per recording session for some channels exceeded 10.

**Figure 2.**
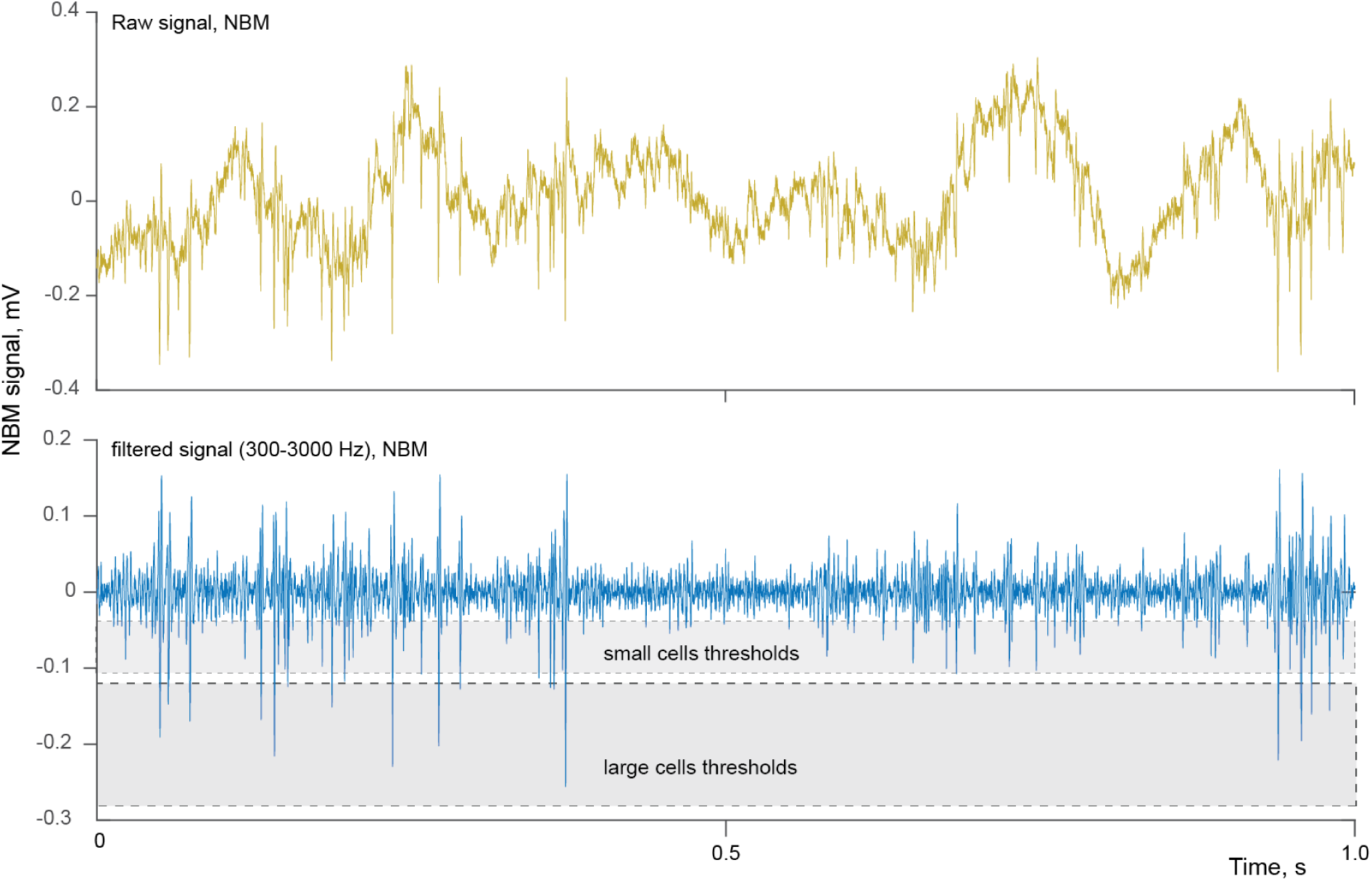
**A second-long example of raw and filtered signal recorded from NBM**. The upper row demonstrates raw NBM signal recorded from one of the animals. The bottom row shows the same signal filtered for spike sorting (bandpass, 300-3000Hz) as well as two thresholding ranges in grey, used to extract signals from two groups of cells, one larger in amplitude and the other smaller.

For each of the identified neurons we calculated multi-taper SFC, separately for every one of the 4 states of vigilance, using the Chronux toolbox (Chronux, 2022; Mitra & Bokil, 2008) and custom-made Matlab scripts. SFC was estimated:

1. Between NBM cells and NBM LFPs (internal NBM coherence)
2. Between cortical cells and NBM LFPs (cortical feedback coherence)

SFC was calculated using 9 orthogonal Slepian tapers and a time bandwidth product of 5 (Mitra and Pesaran 1999; Jarvis and Mitra 2001; Bokil et al. 2007). For data length interval N and frequency bandwidth W, the first K = 2NW − 1 Slepian sequences are concentrated optimally within frequency range [− W, W]. Thus, the minimal frequency range for estimating coherence significance is 2W = (K + 1)/N. We used an interval of N = 2.8 sec, therefore 2W = (9+1)/2.8 = 3.57Hz. Thus, SFC was considered significant if coherence value consistently reached > 95% of Jackknife confidence interval over a frequency range exceeding 3.57Hz.

After doing so for every cell, the resulting coherences per frequency point were transformed into a set of ones and zeros with ones corresponding to significant coherence for that frequency point.

This allowed summing coherences calculated for different cells to find frequency ranges where significant coherence was present for the majority of cells. Finally, peaks of significant coherences at low (<10Hz) and gamma (40-90Hz) ranges were used to create a ***nested*** stimulation pattern for Experiment 2. The nested pattern incorporated both frequencies into bursts of gamma pulses delivered in patches at a lower frequency.

### Experiment 2. Evoked cortical activity by DBS of NBM

Figure 1 panel C schematically shows the shape of the pulse and patterns of stimulation. As mentioned above, we used 3 different patterns of stimulation to evoke cortical responses and conducted stimulations across multiple sleep-wake cycles. For all patterns, individual biphasic pulses lasted 1 ms and 30 pulses were delivered within a single pulse train, with 10 sec interval between trains (10 sec interstimulus interval, ISI, see an example shown by Fig 1 panel E).

Stimulation at frequencies above 100Hz and charge per phase limits above 4 nC/phase are considered unsafe for the nerve tissue (Cogan et al., 2016), therefore both nested and random patterns were limited to <90Hz interval between any 2 pulses and all stimulations conducted with charge per phase not exceeding 0.1 μC.

Pulse train containing 30 pulses lasted 1 sec for nested and random patterns and 1.5 sec for the tonic pattern of 20Hz.

The following patterns of stimulation were studied: nested, tonic (20Hz, as in human clinical trials) and random-generated (as unstructured control).

We started stimulation sessions for each of the patterns from 20 µA per phase current. After collecting enough data for every state of vigilance and every pattern of stimulation, we increased the intensity by 40 µA and repeated the procedure after checking for the appropriate electrode impedance decrease (that commonly rapidly follows stimulation). The intensity was increased in 40 µA steps until the evoked response for every pattern reached saturation, which we defined as significant prolonged decrease of delta activity during SWS - not only immediately after the stimulation, but also at the end of the 10 sec ISI. SWS was chosen as a benchmark of effectiveness for 2 reasons:

1. Unwanted sleepiness and inability to sustain normal level of alertness are important symptoms for most types of dementia (Kasanuki et al., 2018; Elder et al., 2022; Carvalho et al., 2018; Feng et al., 2021), thus it was necessary to establish that stimulation can abolish it in principle;
2. Due to the absence of motion artifacts in SWS and substantial time spent by rodents in that state during the light phase of the cycle, SWS provided the highest amount of good-quality data for the comparative analysis.

The following spectral LFP analysis was done to test (i) whether cortical activity changes in response to stimulation, (ii) which relevant frequency ranges were affected and (iii) whether these responses differed between the patterns of stimulation.

We categorised the states of vigilance and band-pass filtered the data at 2-250 Hz and resampled it at 1000 Hz. LFP spectra were calculated using multi-taper approach, with 5 Slepian tapers and a time-bandwidth product of 3. To avoid transient artifacts caused by the electrical stimulation, we excluded from the analysis a 200 ms interval before the onset of stimulation and a 400 ms interval immediately after the offset of stimulation. To further suppress the artifact, the lower cutoff frequency for bandpass filtering was set at 2Hz.

The results were obtained for each animal at the above-mentioned level of response saturation. The measured value was the integral of spectral density over the frequency range in question.

To compare responses for different patterns within each state of vigilance we used 3 intervals, each 2 secs long: after the stimulation at the beginning of ISI, at the end of the ISI period and background activity without stimulation. The rationale behind this choice was that brain activity disturbance after the stimulation could be long-lasting and not returning to a level normally observed for that state of vigilance by the end of the 10 sec ISI. To detect both the immediate post-stimulus and the long-lasting effects, it was necessary to compare them with the activity without any stimulation (background).

To use SWS as a benchmark activity for detecting effects of stimulation, we always started a recording session from when the animal was asleep and used a 1-2 min period of SWS as the background before starting any stimulation (Fig 1D and 1E). It was not possible to record also REM, AW and QW background levels before each stimulation session because that would have made recording sessions too long and potentially uncomfortable for the animals. Therefore, for these states, we used the background activity recorded during Experiment 1.

Comparisons were made both within each animal and for the entire group. For the comparisons within each animal, we used all individual artifact-free stimulations for the same pattern, intensity level and state of vigilance. The following frequency ranges were used for the analysis at the population level: We were predominantly interested in changes within delta (defined as 2-4.5Hz due to the need to remove low frequency component of a stimulation artifact), theta (5-9Hz), beta-low gamma (20-45Hz) and high gamma (53-80Hz) ranges. However, we noticed broadband changes to cortical activity (see Results for detailed explanation) and therefore included alpha (9-13Hz) and beta (13-20Hz) ranges in the analysis as well.

The reason to break gamma range into 2 parts was the established differences in the relevance of these frequency ranges to AD as well as some other neurological conditions. Early gamma activity deteriorates in AD patients while high gamma changes are less well-known (Casula et al., 2022; Arroyo-García et al., 2021; Murty et al., 2021), while PDD and DBL can be characterised more by general slowing of EEG power (Bonanni et al., 2008; van der Zande et al., 2018), with some evidence for early gamma deterioration (Güntekin et al., 2023). It is therefore important to find out what kind of gamma activity changes occur as a result of NBM DBS. The specific ranges chosen were also dictated by the need to apply a notch filter to the data to remove 50Hz noise in some cases.

In addition, we were interested in the narrow gamma band differences within each individual animal, since highly synchronized neuronal responses can result in narrow-band gamma changes rather than broad-spectrum ones. To analyse that we visualised the frequency spectra and noted all gamma-range differences longer than >3W frequency intervals. The spectra were then compared specifically within these intervals.

For population assessment of the pattern-related effects we treated each cortical electrode as an independent unit of statistics since these site-based results often did not correspond within a single animal. We used Wilcoxon signed-rank test for paired data comparisons of the median values, Wilcoxon rank-sum test for unpaired comparisons and Exact Fisher test for reporting difference in the proportions of the effects between patterns (as categorical variables in contingency tables).

## Results

### Experiment 1. Determining natural frequency peaks of cell-LFP synchronization

#### Spike-field coherence (SFC)

SFC is a normalized index that assesses how well neuronal action potentials (“spikes)” synchronize to continuous LFP oscillation at a particular frequency. The strength of the synchronization is represented by the SFC value, which varies from zero to one. 95% Jackknife confidence intervals for SFC were computed for each cell across the frequency spectrum and separately for each state of vigilance. 9 animals were used for the analysis. Figure 3 provides an example of SFCs obtained for a single NBM cell to NBM LFP at frequencies ranging from 0.5 to 100Hz. Solid lines represent coherence values and dashed lines show 95% confidence interval thresholds.

1. Between NBM cells and NBM LFPs (internal NBM cells coherence)
2. Between cortical cells and NBM LFPs (cortical feedback coherence)

**Figure 3.**
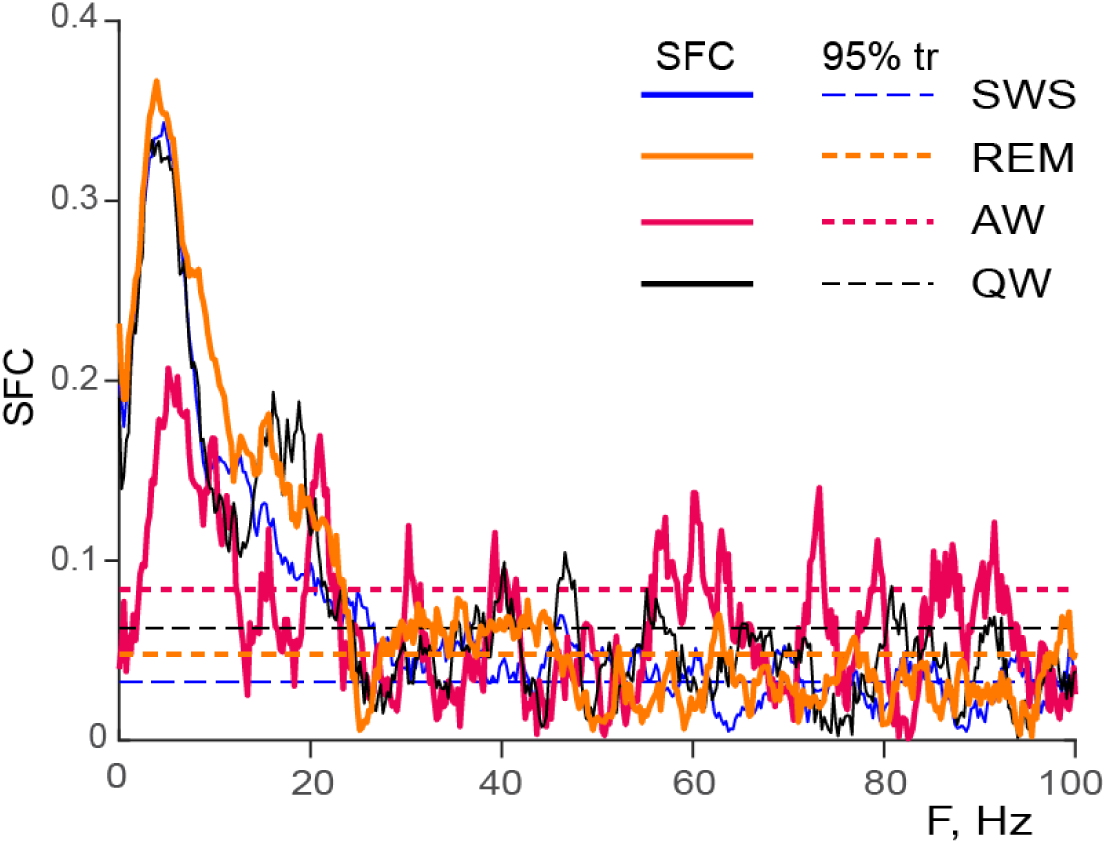
**SFC example for a single NBM cell to NBM LFP**. Solid lines show SFC for each of 4 states of vigilance, while dashed lines represent the corresponding 95% Jackknife confidence intervals (95% tr). SWS is given in blue, REM in orange, AW in red and QW in black. SFC values continuously above threshold for >3.57 Hz frequency interval were considered significant and further analysed by transforming values for significant coherences to ones and for insignificant to zeros, and then summarising resulting values for all cells within 2 groups of coherence measurements:

The significant coherences transformed into cell numbers are shown by figure 3.

##### SFC of NBM cells to NBM LFPs

The upper left panel of Figure 4 shows the sums of significant coherences across 0.5-100Hz frequency range for NBM cells recorded from 2 electrodes in each of 9 animals. The total number of recorded cells was 215. The biggest peak of coherence was in the delta-theta range for all states of vigilance. It was especially prominent and present in 55% of all recorded neurons in SWS, followed by 41% of cells in REM, 35% in QW, and 18% in AW. The second elevation in coherence was observed within 57 - 88 Hz range in SWS, QW and AW, with additional peaks at 28 and 37 Hz for AW, and a REM peak at 88 Hz, likely resulting from REM bursts.

**Figure 4.**
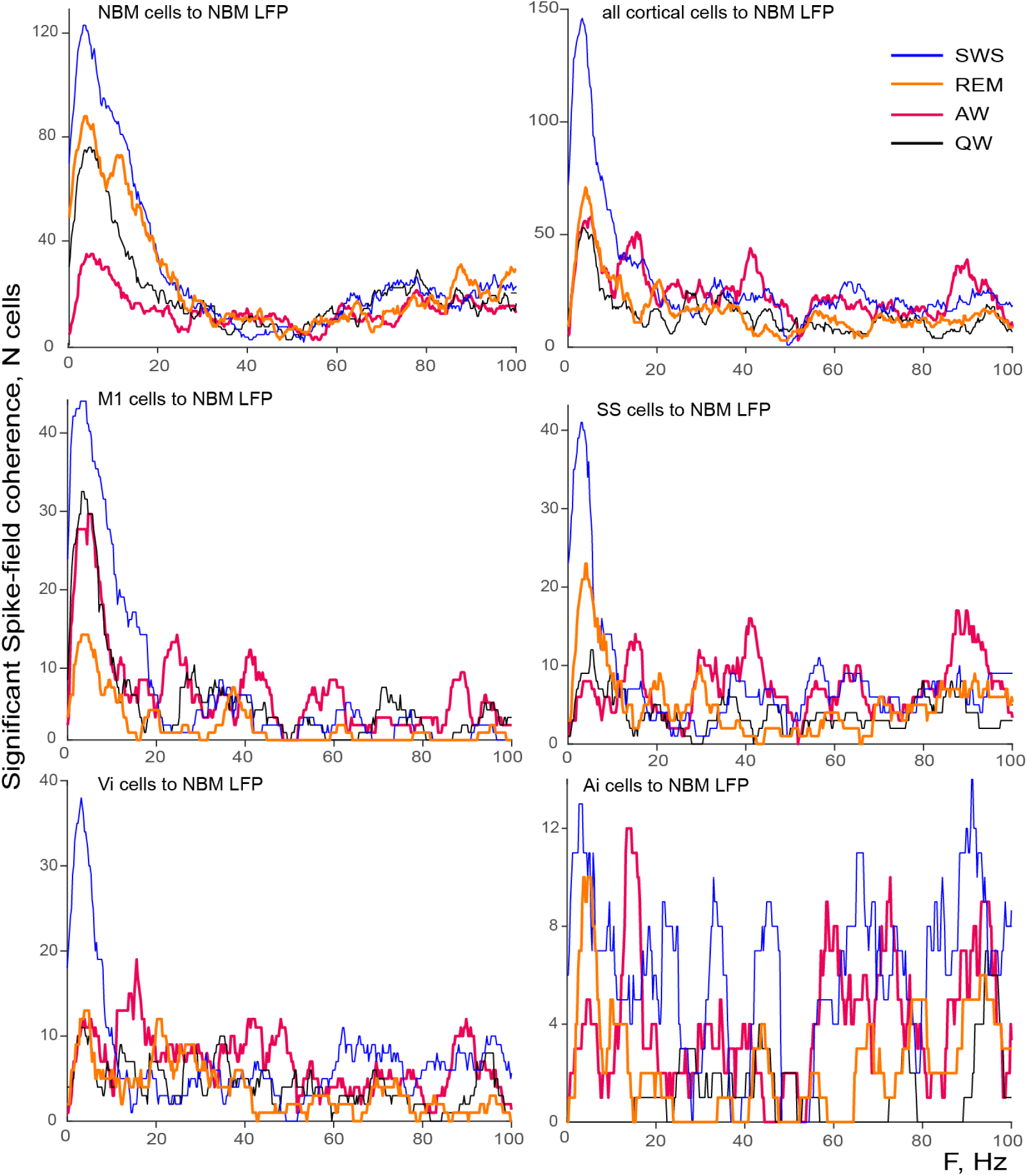
**Cortical neurons which demonstrated significant spike-field coherence to NBM LFP**. The upper row demonstrates SFC numbers for all NBM cells (left panel) and all cortical cells (right panel). The other two rows show SFC numbers for individual cortical areas: primary motor cortex (M1), Supplementary somatosensory (SS), Visceral insula (Vi) and Anterior insula (Ai). Blue lines represent SWS, orange – REM, red – AW and black – QW states of vigilance.

##### SFC of cortical cells to NBM LFPs

Cells recorded from the four cortical areas (M1, SS, Ai and Vi) in 9 animals amounted to 365 neurons, M1 = 110, SS = 92, Ai= 64 and Vi = 99. As can be seen in Figure 4, The most prominent peak registered in the cortex was also in the delta-theta range, especially evident for SWS (40% of all cortical cells) but present in all the other states with the only exception of AW, which had higher peaks in beta and gamma frequency ranges in SS, Vi and Ai. The highest proportion of feedback-synchronizing cells was found in M1, followed by SS and Vi, with the least in Ai.

The results of Experiment 1 informed our decision for the frequencies of the nested frequency pattern. Due to the very prominent presence of low frequencies in both NBM cells to NBM LFP and cortical cells to NBM LFP synchronizations, we used 5Hz as the lower or “nesting” frequency. The higher “nested” frequency for stimulation was chosen for each individual animal as the gamma peak in QW or SWS (whichever was more commonly observed). Individual gamma peaks for the 9 animals were the following: 63, 73, 58, 73, 57, 88, 59, 62, 88 Hz.

### Experiment 2. Evoked cortical activity to NBM DBS

#### Data volume

Among 9 animals successfully implanted with NBM and cortical electrodes, one was excluded from this analysis due to the degradation of one of the NBM electrode’s signal which made bipolar stimulation impossible. For the other 8 animals, stimulations were conducted as both NBM electrodes and 2-4 of the cortical electrodes were preserved by the time of Experiment 2. However, for two of them we could not collect data for tonic stimulation. One mouse developed tonic-clonic seizures specifically resulting from the tonic type of stimulation whereas no seizures were evoked by either nested or random stimulation. The other animal lost NBM signal during tonic stimulation at saturation intensity. Therefore, we present data for 25 nested and random stimulation patterns - evoked responses for 25 individual cortical channels recorded from 8 mice, and 19 data points evoked responses for 19 individual cortical channels recorded from 6 mice – for the tonic stimulation pattern.

#### Responses evoked in different states of vigilance

##### SWS

While in some sessions, delta wave suppression immediately after NBM stimulation could be observed even with naked eye, at the population level it was pronounced and significant (p<0.001) immediately after the stimulation for all cortical channels and all animals even for the lowest intensity of stimulation (20 µA) for each of the 3 stimulation patterns (see Fig 5 for illustration, where the red lines represent Power Spectral Density (PSD), for the ISI interval of 2 sec after the stimulation). PSD is calculated by dividing the power spectrum of a signal by the bandwidth of the frequency components) In all channels, saturation of delta suppression – delta power being below background level by the end of the ISI – was reached at different intensities for different animals. For one animal, the saturation occurred at the lowest intensity of 20 µA, and for another one at the highest intensity of 140 µA. Among the rest, 3 reached saturation at 60 µA and 3 at 100 µA. The suppression of slow wave activity was higher immediately after the stimulation (“start of ISI”) than by the end of ISI (“end of ISI”), as shown by the right panel of Figure 5. This partial recovery of delta activity was also a common and significant effect (p<0.001 for all patterns).

**Fig 5.**
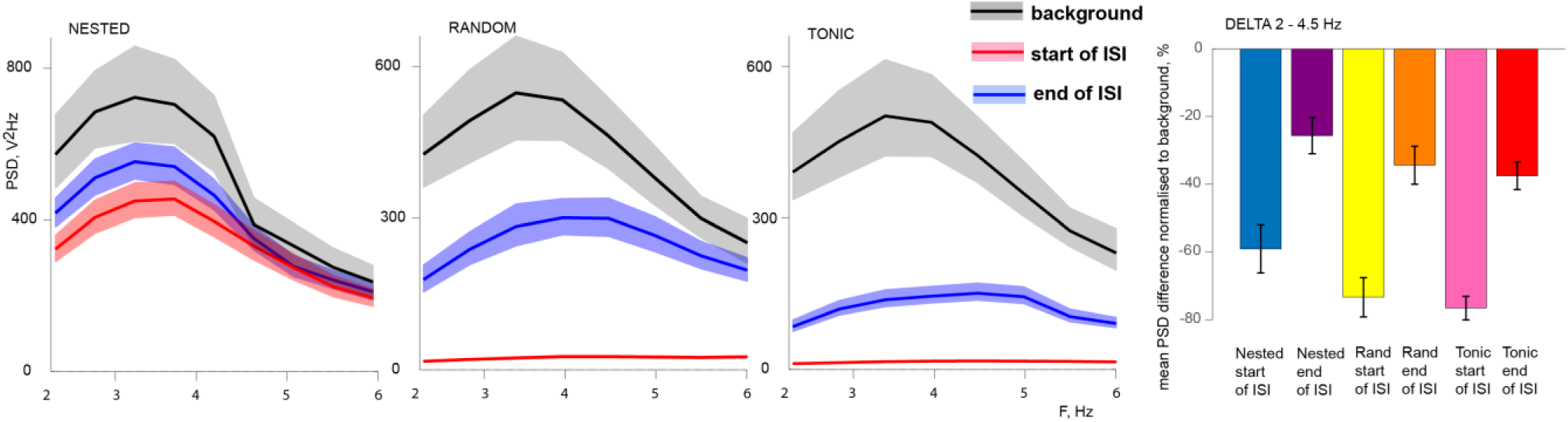
**Examples of delta suppression for a single cortical channel of one mouse and the population response of delta activity to NBM DBS during SWS**. The three panels on the left demonstrate responses for DBS of the same intensity in one cortical channel in one of the animals, shown as Power Spectral Density (PSD). Black solid line demonstrates SWS background delta power, red line shows it for the start of ISI interval and blue for the end of ISI. Light-coloured areas demonstrate standard error of mean. Right panel shows changes in spectral power in percent to background level for the delta range, and standard errors, averaged across all the animals and all cortical channels.

We conclude that suppression of slow wave activity can be reliably achieved by NBM DBS regardless of the pattern of stimulation. In the case of SWS, it is likely to lead to deterioration of sleep quality.

However, as illustrated by Fig 5, delta suppression at the end of ISI was reduced for nested pattern of stimulation. The difference in the degree of suppression was significant (p=0.028) between nested and tonic patterns, while it did not reach significance between nested and random patterns. Although delta suppression remained well below the background level for nested stimulation by the end of ISI, low frequencies recovered faster for this pattern.

Population broad-spectrum high gamma activity was also significantly reduced for all 3 patterns immediately after the stimulation, as well as at the end of the ISI for tonic and nested patterns (p<0.02) (see Fig 6). However, narrow-band high gamma power changes evoked within individual cortical channels demonstrated differences between the patterns of stimulation. Immediately after the stimulation, nested pattern evoked narrow-band enhancement significantly more often than suppression, while tonic and random patterns resulted in more suppression than enhancement (Exact Fisher p= 0.048 and p=0.039 correspondingly, see Table 2). During transitions from sleep to wakefulness (arousal), it is known that there can be a transient suppression of high gamma activity as cortical networks re-engage (Wang et al., 2024).

**Figure 6.**
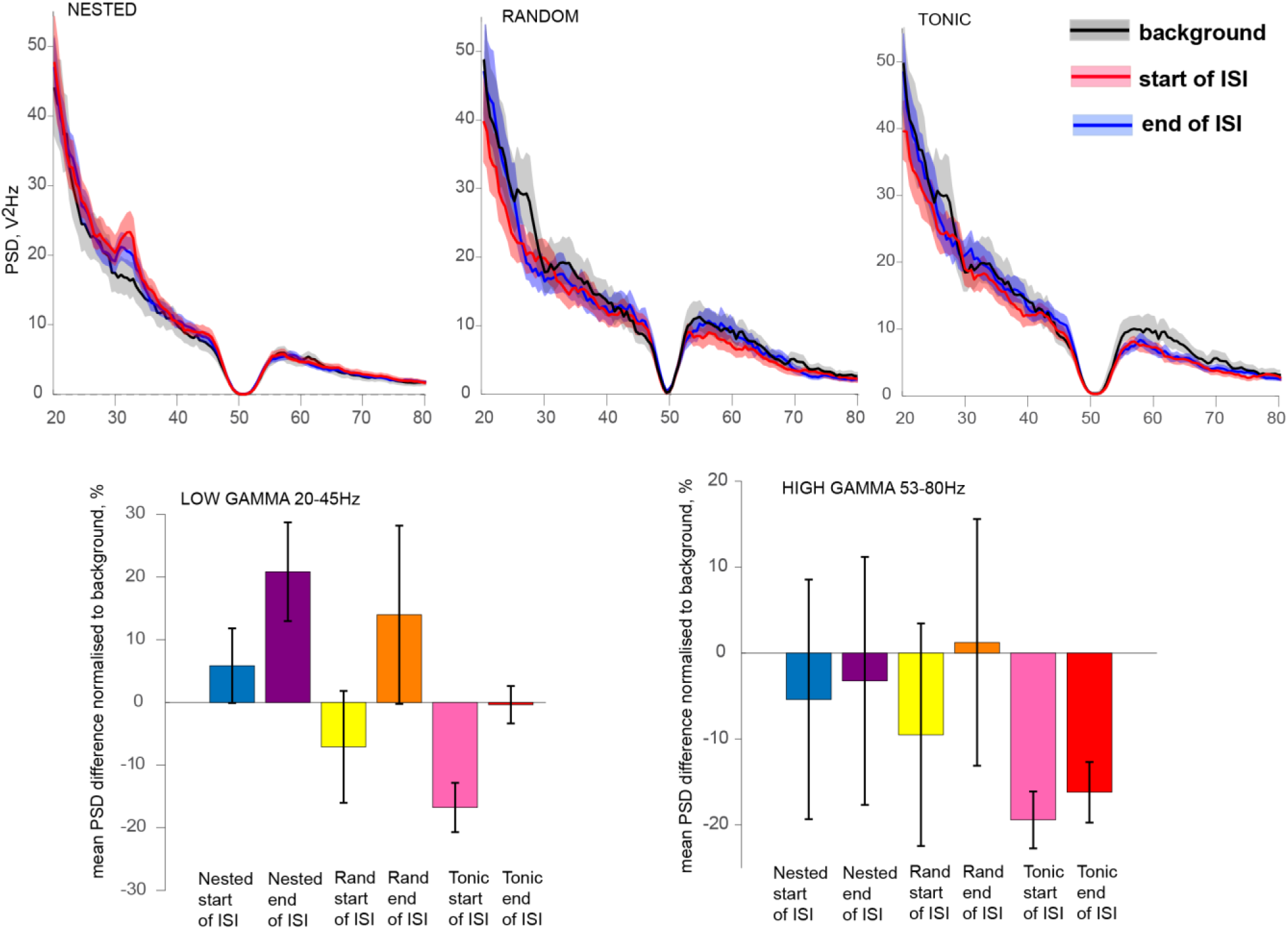
**Examples of gamma responses for a single cortical channel of one mouse and the population response of delta activity to NBM DBS during SWS**. The three top panels demonstrate responses for DBS of the same intensity in one cortical channel in one of the animals. Black solid line demonstrates SWS background gamma power, red line shows it for the start of ISI interval and blue for the end of ISI. Light-coloured areas demonstrate standard error of mean. Bottom panels show changes in spectral power in percent to background level for gamma responses, and standard errors, averaged across all the animals and all cortical channels, for low gamma (left) and high gamma (right) frequency ranges.

**Table 1.** Criteria for categorization of the states of vigilance.

| State of vigilance | Categorization criteria |
| --- | --- |
| Active wakefulness, AW | <5 sec from a movement, desynchronised EEG, gamma activity present, behaviourally active |
| Quiet wakefulness, QW | >5 sec from a movement, no EEG delta activity, behaviourally active |
| Slow wave sleep, SWS | consistent delta waves in EEG, behaviourally inactive |
| REM sleep, REM | consistent theta waves, motion atonia with sporadic jerky movements |

**Table 2.** Types of responses during SWS to stimulation in the high gamma range.

| Stimulation pattern | Beginning of ISI |  | End of ISI |  |
| --- | --- | --- | --- | --- |
|  | enhancement | suppression | enhancement | suppression |
| Nested | 14 | 4 | 12 | 5 |
| Tonic | 4 | 7 | 6 | 6 |
| Random | 6 | 10 | 8 | 13 |

Both the broad-band and narrow-band low gamma activity changes were sensitive to the pattern of stimulation. Nested pattern caused enhancement of the broadband low gamma activity towards the end of ICI (p=0.0087) with no difference to background immediately after the stimulation. This is consistent with the normal awakening response. At the same time, both tonic and random stimulations caused inhibition of broad-band low gamma immediately after the stimulation (p<0.001 for tonic and p=0.03 for random, see Fig 6). The differences in the proportions of the narrow-band enhancement to suppressions between nested and tonic patterns were significant at the beginning of the ISI as well as towards the end of ISI (p=0.0013, p=0.0022). The difference in that proportion between nested and random patterns reached significance towards the end of the ISI (p=0.0028). As seen from Table 3, nested stimulation produced enhancement of the narrow-band cortical activity more often than suppression of it, while that was not the case for the other patterns.

**Table 3.** Types of responses during SWS to stimulation in the early gamma range.

| Stimulation pattern | Beginning of ISI |  | End of ISI |  |
| --- | --- | --- | --- | --- |
|  | enhancement | suppression | enhancement | suppression |
| Nested | 13 | 4 | 24 | 3 |
| Tonic | 1 | 10 | 4 | 7 |
| Random | 6 | 8 | 9 | 11 |

Theta power was also reduced after the stimulation (p<0.02), with the effect lasting towards the end of ISI for tonic and random pattern, but recovering for the nested stimulation. Suppression of activity in the alpha and beta ranges had the same characteristics, as they were suppressed by tonic and random DBS across the entire ISI, while nested pattern caused alpha and beta suppression only immediately after the stimulation, with complete recovery towards the end of the ISI.

##### REM

We report a dramatic reduction of theta activity in response to stimulation delivered during REM sleep. At the level of individual channels, a significant (p<0.05) effect of theta activity decrease was observed for all patterns and 7 animals out of 8, with no noticeable differences between patterns.

Population analysis confirmed the same at p<0.01 for all the patterns and also revealed that the effect was prolonged with theta activity below background level by the end of ISI (see Fig 7). However, it is also essential to note that the effects of stimulation during REM were not limited to theta band. In fact, a significant widespread power decrease was seen in all the bands tested: delta, theta, alpha, beta and low gamma ranges. Only high gamma range seems to be “spared” from the effect, as its power does not change significantly. Although partial recovery of activity was present in all the affected bands towards the end of ISI with power being significantly higher in comparison to the beginning of ISI, this recovery was incomplete and the activity did not reach background level. It seems that the disruption of activity in REM might be the most severe of the detected effects, highlighting the fact that stimulation during REM might be especially counterproductive.

**Figure 7.**
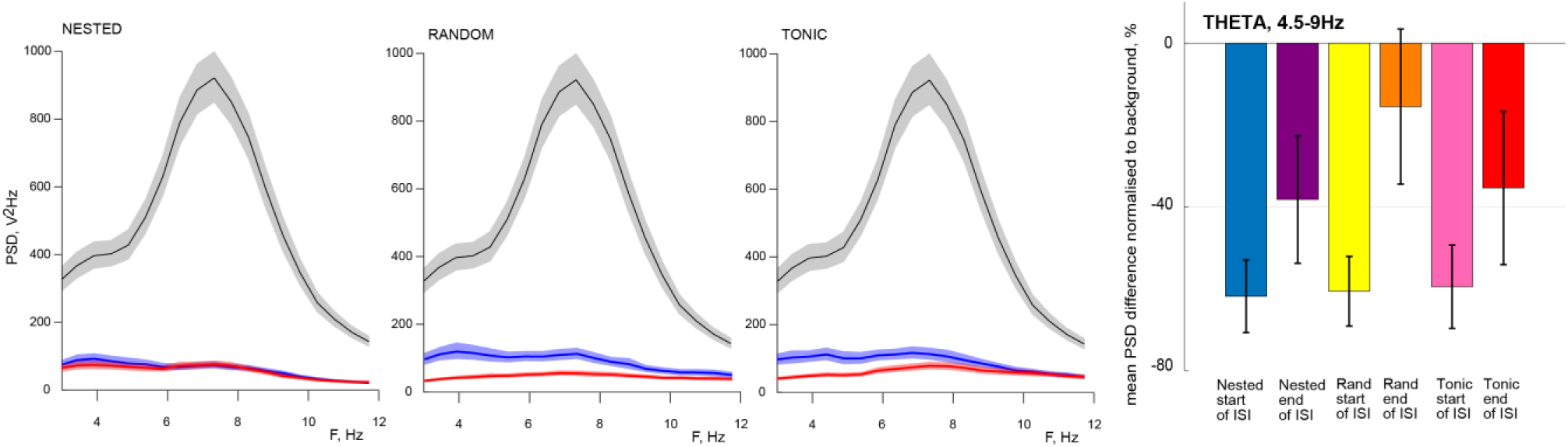
**Examples of theta suppression for a single cortical channel of one mouse and the population response of theta activity to NBM DBS during REM**. The three panels on the left demonstrate responses for DBS of the same intensity in one cortical channel in one of the animals. Black solid line demonstrates SWS background theta power, red line shows it for the start of ISI interval and blue for the end of ISI. Light-coloured areas demonstrate standard error of mean. Right panel shows changes in spectral power in percent to background level for the theta range and standard errors averaged across all the animals and all cortical channels.

##### AW

Although we initially planned to characterise all 4 states of vigilance, we found that stimulation during AW often led to increased movements after stimulation that might have been a desirable behavioural effect showing cholinergic support of actions. However, it also produced motion artifacts, especially in the gamma range. That left us with the only option – to exclude from the analysis all trials containing motion artifacts, which would have reduced statistical power as well as partially defied the purpose of the analysis since the most active states would have been excluded from the AW. Thus, we decided not to present the AW data to avoid this confound.

##### QW

In terms of the possibility to reduce sleepiness and increase alertness by DBS this state is probably the most informative, since the purpose would be to shift the state from the transitional QW to more of the AW-like state.

We found all the patterns to be capable of significantly (p>0.01) reducing delta and theta power after the stimulation, with the effect lasting towards the end of ISI (see illustration for delta power at Figure 8). Thus, sleep reduction by DBS is possible during wakefulness.

**Figure 8.**
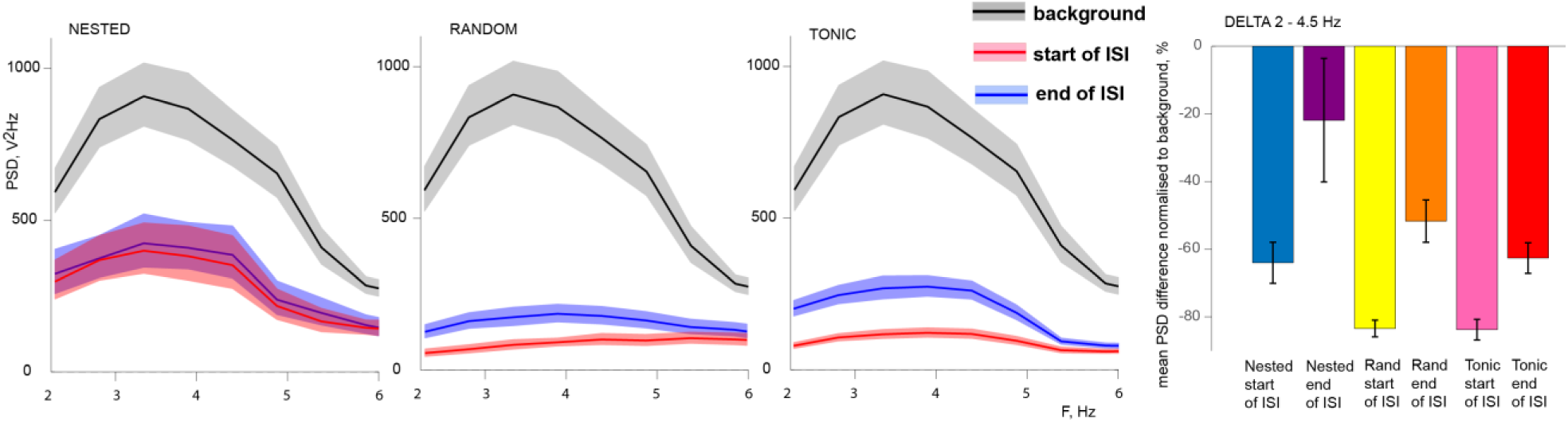
**Examples of delta suppression for a single cortical channel of one mouse and the population response of delta activity to NBM DBS during QW**. The three panels on the left demonstrate responses for DBS of the same intensity in one cortical channel in one of the animals. Black solid line demonstrates QW background delta power, red line shows it for the start of ISI interval and blue for the end of ISI. Light-coloured areas demonstrate standard error of mean. Right panel shows changes in spectral power in percent to background level for the delta range, and standard errors, averaged across all the animals and all cortical channels.

Similar to SWS, delta suppression was least for nested pattern of stimulation. In the case of QW, the differences were significant for both the start of ISI and end of ISI, between nested and tonic patterns (p=0.014, p=0.028) as well as nested and random patterns (p=0.016, p=0.036). Again, delta suppression remained well below the background level for nested stimulation; however, low frequencies were reduced less and probably recovered faster for this pattern (Fig 8).

The effects of DBS in the higher frequency ranges differed between patterns (see Fig 9 for examples and population response). While tonic stimulation severely suppressed activity in the low gamma range immediately after the stimulation (p=0.019) with some recovery towards the end of ISI (p=0.07 for the comparison to background level of low gamma), no such suppression was caused by either nested or random stimulation, with both showing more of a tendency to enhance low gamma activity (Fig 9).

**Figure 9.**
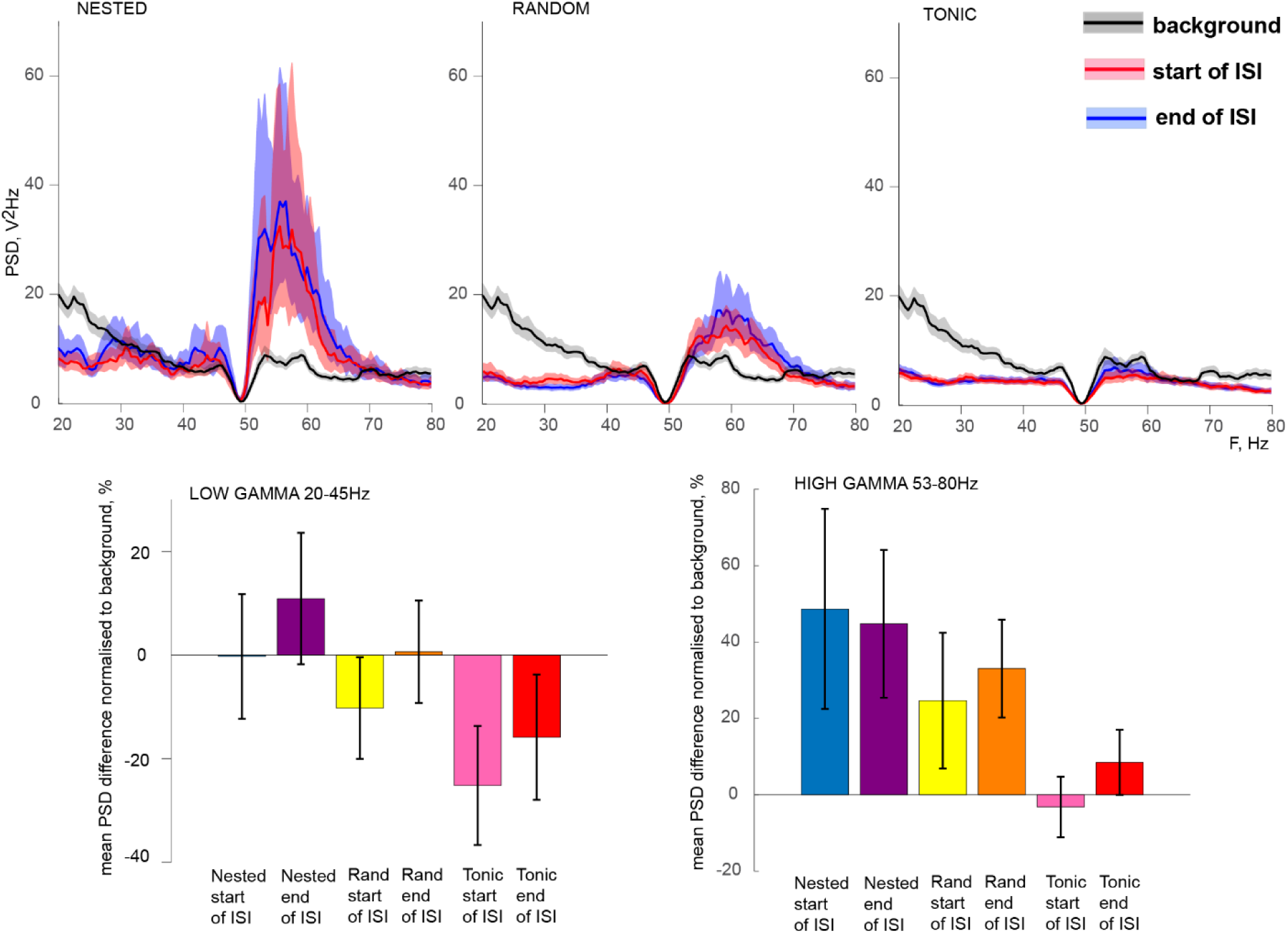
**Examples of gamma responses for a single cortical channel of one mouse and the population response of delta activity to NBM DBS during QW**. The three top panels demonstrate responses for DBS of the same intensity in one cortical channel in one of the animals. Black solid line demonstrates QW background gamma power, red line shows it for the start of ISI interval and blue for the end of ISI. Light-coloured areas demonstrate standard error of mean. Bottom panels show changes in spectral power in percent to background level for gamma responses, and standard errors, averaged across all the animals and all cortical channels, for low gamma (left) and high gamma (right) frequency ranges.

In addition, suppression of activity in the intermediate frequencies – alpha and beta – also demonstrated differences. Both were suppressed by tonic and random DBS across the entire ISI. Nested pattern caused alpha and beta suppression only immediately after the stimulation, with a complete recovery towards the end of the ISI.

We conclude that although all patterns tested were able to suppress sleep-related portions of cortical activity, nested pattern produced effects in the higher frequencies which more closely matched the desired wakefulness-related activity. One obvious result from our study is that stimulation using any pattern should be avoided during both SWS and REM sleep unless the desired effect is to disturb sleep, such as it can be, e.g., in the case of narcolepsy.

## Discussion

The results show that NBM cells tend to synchronize their activities to the local LFP within delta-theta range in all states of vigilance. Similar low-range peak of synchronization occurred between cortical cells and LFP of NBM, indicating that cortical feedback to NBM may also use the same synchronization range, which is not uncommon for long-range connections (Cohen et al., 2021; Myers et al., 2022). This provides grounds for delta-theta frequencies of stimulation bursts - “nesting” frequency that have been suggested from empirical studies (e.g., reviewed by Kumbhare et al., 2018). The highest peak of gamma synchronization of NBM cells was observed between 57 and 88 Hz in SWS, QW and AW, with additional peaks at 28 and 37 Hz for AW, and a REM peak at 88 Hz.

Cholinergic input from NBM activates somatostatin (SOM) interneurons and inhibits parvalbumin PV interneurons in the cortex (Kruglikov & Rudy, 2008; Fanselow et al., 2008; Poorthuis et al., 2014). In turn, SOM interneurons promote beta-low gamma oscillations in the cortex, while PV cells produce high gamma responses (Chen et al., 2017). The effects of stimulation we observed in the gamma range were in line with the increase of the cholinergic influence of NBM. It has also been established that stimulation of different NBM cells may result in dramatically different outcomes for the cortical activity and these types of cells have dissimilar spike rates (Xu et al., 2015; Anaclet et al., 2015; Kim et al., 2015). Therefore, quite possibly, they can be activated or inhibited by different stimulation frequencies or patterns. Since the goal of Experiment 2 was to enhance activity in NBM and reduce sleepiness, we chose 5 Hz and individual QW and SWS gamma peaks (which fell between 57 and 88 Hz in all animals). This approach turned out to be the only one that supported beta-low gamma activity after the stimulation delivered in SWS, as well as suppressing it during QW. While all patterns reliably suppressed low frequency activity and therefore can be useful in reducing EDS, they can lead to unwanted decrease of gamma-range activity, especially for the tonic type of stimulation. This effect of decrease of low-gamma power can be attributed to inhibition of basal forebrain parvalbumin neurons since they have been shown to enhance cortical gamma band oscillations when optogenetically stimulated, with their optogenetic inhibition leading to gamma power reduction (Kim et al., 2015). These cells were reported to follow optical stimulation at 40-60Hz; however, electrical stimulation is less selective and can, e.g., overstimulate local inhibitory circuits, inhibiting certain types of neuronal outputs of NBM. It seems that the choice of tonic mode of stimulation for human clinical work might have been an unfortunate one.

A robust suppression of low frequency power in the cortex was induced by all examined patterns in all states of vigilance examined. This result has two important consequences. On the one hand, NBM DBS is likely to be effective in preventing sleepiness in various conditions that have EDS as part of their pathology, which includes AD, PDD and DLB (Kasanuki et al., 2018; Elder et al., 2022; Carvalho et al., 2018; Feng et al., 2021). On the other hand, it can also possibly be extended to pharmacoresistant cases of narcolepsy or hypersomnia, which has been estimated at above 10% of such patients (Anderson et al., 2007; Wise et al., 2007). It is also critical to provide NBM stimulation strictly during the time of desired wakefulness to avoid exacerbation of sleep disturbances that constitute an important part of dementia pathology as well (Boeve et al., 1998; McKeith et al., 2017; Liguori et al., 2014; Mander et al., 2015; Kuang et al., 2021; Menza, 2010; Zhang et al., 2022; Minakawa, 2022). We found that NBM DBS has the potential to disrupt both SWS and REM sleep, irrespective of the type of stimulation and an exceptionally broad-band decrease of cortical activity when it was applied in REM. This is consistent with the suggestion that continuous DBS across sleep-wake cycle can be disruptive rather than helpful (reviewed by Nazmuddin et al., 2021; Kumbhare et al., 2018).

One of the observed characteristics of the cortical response to DBS was a trade-off between the depth of delta suppression and the ability to support gamma frequencies. The effect of broad-band suppression of cortical activity is unusual by itself, with the nearest similarity to that being the effect of psychedelic drugs (Vejmola et al., 2021). Our work cannot provide a direct answer to why such broad-band suppression happens and whether it is indeed similar to the perceptual effects of psychedelics. However, we can see that the effect is likely to be associated with the tonic type of stimulation and to a lesser extent with random stimulation as well. Unlike nested pattern that utilized theta and high gamma frequencies, random pattern by nature also contains intermediate frequencies closer to the tonic 20Hz frequency. Considering the therapeutic potential of psychedelic drugs (De Gregorio et al. 2021), it would be interesting to assess whether tonic NBM DBS can produce similar effects.

We suggest below some further directions that could be useful for improving the method of NBM DBS.

Although human clinical trials are obviously the ultimate way to check whether the DBS helps real patients, some animal experiments can help to clarify the situation before proceeding to clinical trials. Firstly, animal experiments would allow choosing the conditions that could benefit from NBM DBS. Although small-size clinical trials were done for both PDD and AD, the effects of stimulation might not necessarily be equally helpful for both, especially if designed in exactly the same way. AD can be characterised by a reduction of low gamma activity (Casula et al., 2022; Arroyo-García et al., 2021; Murty et al., 2021) and thus can be more responsive to a type of stimulation that helps to preserve it, which is less common for PD. In addition, DBS is already used in PD to alleviate motion-related symptoms, sometimes provoking cognitive decline (Morrison et al., 2004; Hariz et al., 2008; Rothlind et al., 2015). Thus, in PDD there may be a need to conduct two types of stimulation, one of the subthalamic area to help with movements and the other of NBM to counteract cognitive decline, similar to the successful dual-target approach of Freund et al. (2009). This would necessitate a different design of the DBS system with different requirements on the battery capacity and, possibly, different intensity of stimulation in AD. At the same time, ESD is most severe in DLB (Elder et al., 2022), thus EDS-targeting approach can be most fruitful in DLB. Animal experiments involving various models of diseases such as AD, PDD, DLB and narcolepsy can provide information on the most promising targets for clinical trials to focus on. Perhaps, it would also be useful to include a model of major depressive disorder (MDD) in the animal studies to assess whether this “psychedelic-like” effect has any therapeutic potential. Animal studies can also clarify whether stimulation can still be effective despite the loss of cholinergic cells in NBM, and verify whether cortical decrease of slow wave activity and increase in gamma power can be viewed as a biomarker of DBS success.

It seems essential to use intermittent nested stimulation that includes both low and gamma frequencies to stimulate NBM in a way that not only reduces slow wave activity, but allows gamma-range activity in the cortex to emerge once EDS is suppressed. Perhaps even more important would be to develop closed-loop DBS systems capable of turning off upon the onset of sleep and/or when delta activity in the cortex increases around the desired bedtime, preventing from stimulation during SWS or REM sleep. This addition to DBS system can be made by recording EEG and adding a classifier based on EEG power bands and their relationships to distinguish sleep from wakefulness (Lambert and Peter-Derex, 2023). This system would also allow correlating the effectiveness of stimulation to the evoked changes in EEG and constantly adjusting stimulation parameters to the severity of EDS. Alternatively, in a simplified version of the system, DBS can be pre-set in accordance to a patient’s sleep-wake cycle, with stimulation that starts at a particular time of the day and gets switched off at some point before bedtime.

## Notes

### Competing Interest Statement

The authors have declared no competing interest.

## References

Akam, T., & Kullmann, D. M. (2014). Oscillatory multiplexing of population codes for selective communication in the mammalian brain. Nature Reviews Neuroscience, 15(2), 111–122. 10.1038/nrn3668

Anaclet, C., Pedersen, N. P., Ferrari, L. L., Venner, A., Bass, C. E., Arrigoni, E., & Fuller, P. M. (2015). Basal forebrain control of wakefulness and cortical rhythms. Nature Communications, 6, Article 8744. 10.1038/ncomms9744

Anderson, E. B., Mitchell, J. F., & Reynolds, J. H. (2011). Attentional modulation of firing rate varies with burstiness across putative pyramidal neurons in macaque visual area V4. Journal of Neuroscience, 31(30), 10983–10992. 10.1523/JNEUROSCI.0027-11.2011

Anderson, K. N., Pilsworth, S., Sharples, L. D., Smith, I. E., & Shneerson, J. M. (2007). Idiopathic hypersomnia: A study of 77 cases. Sleep, 30(10), 1274–1281. 10.1093/sleep/30.10.1274

Arnold, S. E., Hyman, B. T., Flory, J., Damasio, A. R., & Van Hoesen, G. W. (1991). The topographical and neuroanatomical distribution of neurofibrillary tangles and neuritic plaques in the cerebral cortex of patients with Alzheimer’s disease. Cerebral Cortex, 1(1), 103–116. 10.1093/cercor/1.1.103

Arroyo-García, L. E., Isla, A. G., Andrade-Talavera, Y., Balleza-Tapia, H., Loera-Valencia, R., Alvarez-Jimenez, L., Pizzirusso, G., Tambaro, S., Nilsson, P., & Fisahn, A. (2021). Impaired spike-gamma coupling of area CA3 fast-spiking interneurons as the earliest functional impairment in the AppNL-G-F mouse model of Alzheimer’s disease. Molecular Psychiatry, 26, 5557–5567. 10.1038/s41380-021-01232-9

Bastos, A. M., Vezoli, J., Bosman, C. A., Schoffelen, J.-M., Oostenveld, R., Dowdall, J. R., De Weerd, P., Kennedy, H., & Fries, P. (2015). Visual areas exert feedforward and feedback influences through distinct frequency channels. Neuron, 85(2), 390–401. 10.1016/j.neuron.2014.12.018

Boeve, B. F., Silber, M. H., Ferman, T. J., Kokmen, E., Smith, G. E., Ivnik, R. J., Parisi, J. E., Olson, E. J., & Petersen, R. C. (1998). REM sleep behavior disorder and degenerative dementia: An association likely reflecting Lewy body disease. Neurology, 51(2), 363–370. 10.1212/wnl.51.2.363

Bonanni, L., Thomas, A., Tiraboschi, P., Perfetti, B., Varanese, S., & Onofrj, M. (2008). EEG comparisons in early Alzheimer’s disease, dementia with Lewy bodies and Parkinson’s disease with dementia patients with a 2-year follow-up. Brain, 131(3), 690–705. 10.1093/brain/awm322

Buschman, T. J., & Miller, E. K. (2007). Top-down versus bottom-up control of attention in the prefrontal and posterior parietal cortices. Science, 315(5820), 1860–1862. 10.1126/science.1138071

Buzsáki, G. (2002). Theta oscillations in the hippocampus. Neuron, 33(3), 325–340. 10.1016/S0896-6273(02)00586-X

Buzsáki, G., & Wang, X.-J. (2012). Mechanisms of gamma oscillations. Annual Review of Neuroscience, 35, 203–225. 10.1146/annurev-neuro-062111-150444

Carvalho, D. Z., St Louis, E. K., Knopman, D. S., Boeve, B. F., Lowe, V. J., Roberts, R. O., Mielke, M. M., Przybelski, S. A., Machulda, M. M., Petersen, R. C., Jack, C. R. Jr., & Vemuri, P. (2018). Association of excessive daytime sleepiness with longitudinal β-amyloid accumulation in elderly persons without dementia. JAMA Neurology, 75(6), 672–680. 10.1001/jamaneurol.2018.0049

Casula, E. P., Pellicciari, M. C., Bonnì, S., Borghi, I., Maiella, M., Assogna, M., Minei, M., Motta, C., D’Acunto, A., Porrazzini, F., Pezzopane, V., Mencarelli, L., Roncaioli, A., Rocchi, L., Spampinato, D. A., Caltagirone, C., Santarnecchi, E., Martorana, A., & Koch, G. (2022). Decreased frontal gamma activity in Alzheimer disease patients. Annals of Neurology, 92(3), 464–475. 10.1002/ana.26444

Chang, H., Tang, W., Wulf, A. M., Nyasulu, T., Wolf, M. E., Fernandez-Ruiz, A., & Oliva, A. (2025). Sleep microstructure organizes memory replay. Nature, 637, 1161–1169. 10.1038/s41586-024-XXXXX

Chaves-Coira, I., Martín-Cortecero, J., Nuñez, A., & Rodrigo-Angulo, M. L. (2018). Basal forebrain nuclei display distinct projecting pathways and functional circuits to sensory primary and prefrontal cortices in the rat. Frontiers in Neuroanatomy, 12, Article 69. 10.3389/fnana.2018.00069

Chen, G., Zhang, Y., Li, X., … Tao, H. W., Rasch, M. J., & Zhang, X. (2017). Distinct inhibitory circuits orchestrate cortical beta and gamma band oscillations. Neuron, 96(6), 1403–1418.e6. 10.1016/j.neuron.2017.11.033

Cogan, S. F., Ludwig, K. A., Welle, C. G., & Takmakov, P. (2016). Tissue damage thresholds during therapeutic electrical stimulation. Journal of Neural Engineering, 13(2), 021001. 10.1088/1741-2560/13/2/021001

Cohen, M. X., Englitz, B., & França, A. S. C. (2021). Large-scale and multiscale networks in the rodent brain during novelty exploration. eNeuro, 8(3), ENEURO.0494-20.2021. 10.1523/ENEURO.0494-20.2021

De Gregorio, D., Aguilar-Valles, A., Preller, K. H., Heifets, B. D., Hibicke, M., Mitchell, J., & Gobbi, G. (2021). Hallucinogens in mental health: Preclinical and clinical studies on LSD, psilocybin, MDMA, and ketamine. Journal of Neuroscience, 41(5), 891–900. 10.1523/JNEUROSCI.1659-20.2020

Dong, H. W. (2008). The Allen reference atlas (Book + CD-ROM): A digital color brain atlas of the C57BL/6J male mouse. Wiley.

Elder, G. J., Lazar, A. S., Alfonso-Miller, P., & Taylor, J. P. (2022). Sleep disturbances in Lewy body dementia: A systematic review. International Journal of Geriatric Psychiatry, 37(10), Article e5814. 10.1002/gps.5814

Fanselow, E. E., Richardson, K. A., & Connors, B. W. (2008). Selective, state-dependent activation of somatostatin-expressing inhibitory interneurons in mouse neocortex. Journal of Neurophysiology, 100, 2640–2652. 10.1152/jn.90691.2008

Feng, F., Cai, Y., Hou, Y., Ou, R., Jiang, Z., & Shang, H. (2021). Excessive daytime sleepiness in Parkinson’s disease: A systematic review and meta-analysis. Parkinsonism & Related Disorders, 85, 133–140. 10.1016/j.parkreldis.2021.02.016

Ferreri, F., Pauri, F., Pasqualetti, P., Fini, R., Dal Forno, G., & Rossini, P. M. (2003). Motor cortex excitability in Alzheimer’s disease: A transcranial magnetic stimulation study. Annals of Neurology, 53(1), 102–108. 10.1002/ana.10416

Fibiger, H. C. (1982). The organization and some projections of cholinergic neurons of the mammalian forebrain. Brain Research Reviews, 4(3), 327–388. 10.1016/0165-0173(82)90011-x

Freund, H.-J., Kuhn, J., Lenartz, D., Mai, J. K., Schnell, T., Klosterkoetter, J., & Sturm, V. (2009). Cognitive functions in a patient with Parkinson-dementia syndrome undergoing deep brain stimulation. Archives of Neurology, 66(6), 781–785. 10.1001/archneurol.2009.102

Fries, P., Roelfsema, P. R., Engel, A. K., König, P., & Singer, W. (1997). Synchronization of oscillatory responses in visual cortex correlates with perception in interocular rivalry. Nature, 389(6650), 251–254. 10.1038/38446

Fries, P. (2005). A mechanism for cognitive dynamics: Neuronal communication through neuronal coherence. Trends in Cognitive Sciences, 9(10), 474–480. 10.1016/j.tics.2005.08.011

Gratwicke, J., Kahan, J., Zrinzo, L., Hariz, M., Limousin, P., Foltynie, T., & Jahanshahi, M. (2013). The nucleus basalis of Meynert: A new target for deep brain stimulation in dementia? Neuroscience & Biobehavioral Reviews, 37(10 Pt 2), 2676–2688. 10.1016/j.neubiorev.2013.09.003

Gratwicke, J., Zrinzo, L., Kahan, J., Peters, A., Beigi, M., Akram, H., Hyam, J., Oswal, A., Day, B., Mancini, L., Thornton, J., Yousry, T., Limousin, P., Hariz, M., Jahanshahi, M., & Foltynie, T. (2018). Bilateral deep brain stimulation of the nucleus basalis of Meynert for Parkinson disease dementia: A randomized clinical trial. JAMA Neurology, 75(2), 169–178. 10.1001/jamaneurol.2017.3762

Gray, C. M., König, P., Engel, A. K., & Singer, W. (1989). Oscillatory responses in cat visual cortex exhibit inter-columnar synchronization which reflects global stimulus properties. Nature, 338, 334–337. 10.1038/338334a0

Güntekin, B., Bölükbaş, B., Duygun, R., Kuloğlu, H. B., Yemeniciler, İ., Yıldırım, E., Aktürk, T., & Yener, G. (2023). How do event-related gamma oscillations differ in Alzheimer’s disease from other types of dementia? Biomarkers. Advance online publication. 10.1002/alz.072960

Hariz, M. I., Rehncrona, S., Quinn, N. P., Speelman, J. D., & Wensing, C.; Multicentre Advanced Parkinson’s Disease Deep Brain Stimulation Group. (2008). Multicenter study on deep brain stimulation in Parkinson’s disease: An independent assessment of reported adverse events at 4 years. Movement Disorders, 23(3), 416–421. 10.1002/mds.21888

Irmak, S. O., & de Lecea, L. (2014). Basal forebrain cholinergic modulation of sleep transitions. Sleep, 37(12), 1941–1951. 10.5665/sleep.4246

Jarvis, M. R., & Mitra, P. P. (2001). Sampling properties of the spectrum and coherency of sequences of action potentials. Neural Computation, 13, 717–749.

Kasanuki, K., Ferman, T. J., Murray, M. E., Heckman, M. G., Pedraza, O., Hanna Al-Shaikh, F. S., Mishima, T., Diehl, N. N., van Gerpen, J. A., Uitti, R. J., Wszolek, Z. K., Graff-Radford, N. R., & Dickson, D. W. (2018). Daytime sleepiness in dementia with Lewy bodies is associated with neuronal depletion of the nucleus basalis of Meynert. Parkinsonism & Related Disorders, 50, 99–103. 10.1016/j.parkreldis.2018.02.003

Kim, T., Thankachan, S., McKenna, J. T., McNally, J. M., Yang, C., Choi, J. H., Chen, L., Kocsis, B., Deisseroth, K., Strecker, R. E., Basheer, R., Brown, R. E., & McCarley, R. W. (2015). Cortically projecting basal forebrain parvalbumin neurons regulate cortical gamma band oscillations. Proceedings of the National Academy of Sciences of the United States of America, 112(11), 3535–3540. 10.1073/pnas.1413625112

Kitamura, J., Nagai, M., Ueno, H., Ohshita, T., Kikumoto, M., Toko, M., Kato, M., Dote, K., Yamashita, H., & Kario, K. (2020). The insular cortex, Alzheimer disease pathology, and their effects on blood pressure variability. Alzheimer Disease & Associated Disorders, 34(3), 282–291. 10.1097/WAD.0000000000000340

Klinzing, J. G., Niethard, N., & Born, J. (2019). Mechanisms of systems memory consolidation during sleep. Nature Neuroscience, 22, 1598–1610. 10.1038/s41593-019-0467-3

Koulousakis, P., van den Hove, D., Visser-Vandewalle, V., & Sesia, T. (2020). Cognitive improvements after intermittent deep brain stimulation of the nucleus basalis of Meynert in a transgenic rat model for Alzheimer’s disease: A preliminary approach. Journal of Alzheimer’s Disease, 73, 461–466. 10.3233/JAD-190919

Kruglikov, I., & Rudy, B. (2008). Perisomatic GABA release and thalamocortical integration onto neocortical excitatory cells are regulated by neuromodulators. Neuron, 58(6), 911–924. 10.1016/j.neuron.2008.04.024

Kuang, H., Zhu, Y. G., Zhou, Z. F., Yang, M. W., Hong, F. F., & Yang, S. L. (2021). Sleep disorders in Alzheimer’s disease: The predictive roles and potential mechanisms. Neural Regeneration Research, 16(10), 1965–1972. 10.4103/1673-5374.308071

Kuhn, J., Hardenacke, K., Lenartz, D., Gruendler, T., Ullsperger, M., Bartsch, C., Mai, J. K., Zilles, K., Bauer, A., Matusch, A., Schulz, R.-J., Noreik, M., Bührle, C. P., Maintz, D., Woopen, C., Häussermann, P., Hellmich, M., Klosterkötter, J., Wiltfang, J., … Sturm, V. (2015). Deep brain stimulation of the nucleus basalis of Meynert in Alzheimer’s dementia. Molecular Psychiatry, 20(3), 353–360. 10.1038/mp.2014.32

Kumbhare, D., Palys, V., Toms, J., Wickramasinghe, C. S., Amarasinghe, K., Manic, M., Hughes, E., & Holloway, K. L. (2018). Nucleus Basalis of Meynert stimulation for dementia: Theoretical and technical considerations. Frontiers in Neuroscience, 12, 614. 10.3389/fnins.2018.00614

Kumbhare, D., Rajagopal, M., Toms, J., Freelin, A., Weistroffer, G., McComb, N., Karnam, S., Azghadi, A., Murnane, K. S., Baron, M. S., & Holloway, K. L. (2024). Deep brain stimulation of nucleus basalis of Meynert improves learning in rat model of dementia. bioRxiv [Preprint]. 10.1101/2024.04.05.588271

Lambert, I., & Peter-Derex, L. (2023). Spotlight on sleep stage classification based on EEG. Nature and Science of Sleep, 15, 479–490. 10.2147/NSS.S401270

Lee, D. J., Milosevic, L., Gramer, R., Sasikumar, S., Al-Ozzi, T. M., De Vloo, P., Dallapiazza, R. F., Elias, G. J. B., Cohn, M., Kalia, S. K., Hutchison, W. D., Fasano, A., & Lozano, A. M. (2020). Nucleus basalis of Meynert neuronal activity in Parkinson’s disease. Journal of Neurosurgery, 132(2), 574–582. 10.3171/2018.11.JNS182386

Lee, J. E., Jeong, D. U., Lee, J., Chang, W. S., & Chang, J. W. (2016). The effect of nucleus basalis magnocellularis deep brain stimulation on memory function in a rat model of dementia. BMC Neurology, 16, 6. 10.1186/s12883-016-0529-z

Lee, M. G., Hassani, O. K., Alonso, A., & Jones, B. E. (2005). Cholinergic basal forebrain neurons burst with theta during waking and paradoxical sleep. Journal of Neuroscience, 25(17), 4365–4369. 10.1523/JNEUROSCI.0178-05.2005

Liguori, C., Romigi, A., Nuccetelli, M., Zannino, S., Sancesario, G., Martorana, A., Albanese, M., Mercuri, N. B., Izzi, F., Bernardini, S., Nitti, A., Sancesario, G. M., Sica, F., Marciani, M. G., & Placidi, F. (2014). Orexinergic system dysregulation, sleep impairment, and cognitive decline in Alzheimer disease. JAMA Neurology, 71(12), 1498–1500. 10.1001/jamaneurol.2014.2510

Lin, S.-C., & Nicolelis, M. A. L. (2008). Neuronal ensemble bursting in the basal forebrain encodes salience irrespective of valence. Neuron, 59(1), 138–149. 10.1016/j.neuron.2008.04.031

Liu, A. K., Chang, R. C. C., Pearce, R. K. B., & Gentleman, S. M. (2015). Nucleus basalis of Meynert revisited: Anatomy, history and differential involvement in Alzheimer’s and Parkinson’s disease. Acta Neuropathologica, 129, 527–540. 10.1007/s00401-015-1392-5

Liu, H., Temel, Y., Boonstra, J., & Hescham, S. (2020). The effect of fornix deep brain stimulation in brain diseases. Cellular and Molecular Life Sciences, 77(17), 3279–3291. 10.1007/s00018-020-03456-4

Liu, R., Crawford, J., Callahan, P. M., Terry, A. V. Jr., Constantinidis, C., & Blake, D. T. (2017). Intermittent stimulation of the nucleus basalis of Meynert improves working memory in adult monkeys. Current Biology, 27(17), 2640–2646.e4. 10.1016/j.cub.2017.07.021

Long, S., Ding, R., Wang, J., Yu, Y., Lu, J., & Yao, D. (2021). Sleep quality and electroencephalogram delta power. Frontiers in Neuroscience, 15, Article 803507. 10.3389/fnins.2021.803507

Lundqvist, M., Rose, J., Brincat, S. L., Warden, M. R., Buschman, T. J., Herman, P., & Miller, E. K. (2022). Reduced variability of bursting activity during working memory. Scientific Reports, 12, 15050. 10.1038/s41598-022-18577-y

Mander, B. A., Marks, S. M., Vogel, J. W., Rao, V., Lu, B., Saletin, J. M., Ancoli-Israel, S., Jagust, W. J., & Walker, M. P. (2015). β-amyloid disrupts human NREM slow waves and related hippocampus-dependent memory consolidation. Nature Neuroscience, 18, 1051–1057.

McKeith, I. G., Boeve, B. F., Dickson, D. W., Halliday, G., Taylor, J.-P., Weintraub, D., Aarsland, D., Galvin, J., Attems, J., Ballard, C. G., … Kosaka, K. (2017). Diagnosis and management of dementia with Lewy bodies: Fourth consensus report of the DLB Consortium. Neurology, 89(1), 88–100. 10.1212/WNL.0000000000004058

Menza, M., DeFronzo Dobkin, R., Marin, H., & Bienfait, K. (2010). Sleep disturbances in Parkinson’s disease [Unpublished manuscript]. National Institutes of Health. https://www.ncbi.nlm.nih.gov/pmc/articles/PMC2840057/

Mesulam, M. M., & Geula, C. (1988). Nucleus basalis (Ch4) and cortical cholinergic innervation in the human brain: Observations based on the distribution of acetylcholinesterase and choline acetyltransferase. Journal of Comparative Neurology, 275, 216–240. 10.1002/cne.902750205

Mesulam, M. M., Mufson, E. J., Levey, A. I., & Wainer, B. H. (1983). Cholinergic innervation of cortex by the basal forebrain: Cytochemistry and cortical connections of the septal area, diagonal band nuclei, nucleus basalis (substantia innominata), and hypothalamus in the rhesus monkey. Journal of Comparative Neurology, 214, 170–197. 10.1002/cne.902140206

Minakawa, E. N. (2022). Bidirectional relationship between sleep disturbances and Parkinson’s disease. Frontiers in Neurology, 13, Article 927994. 10.3389/fneur.2022.927994

Mitra, P. P., & Bokil, H. (2008). Observed brain dynamics. Oxford University Press.

Mitra, P. P., & Pesaran, B. (1999). Analysis of dynamic brain imaging data. Biophysical Journal, 76, 691–708.

Morrison, C. E., Borod, J. C., Perrine, K., Beric, A., Brin, M. F., Rezai, A., et al. (2004). Neuropsychological functioning following bilateral subthalamic nucleus stimulation in Parkinson’s disease. Archives of Clinical Neuropsychology, 19(2), 165–181. 10.1016/S0887-6177(03)00071-9

Murty, D. V. P. S., Manikandan, K., Kumar, W. S., Ramesh, R. G., Purokayastha, S., Nagendra, B., ML, A., Balakrishnan, A., Javali, M., & Ray, S. (2021). Stimulus-induced gamma rhythms are weaker in human elderly with mild cognitive impairment and Alzheimer’s disease. eLife, 10, e61666. 10.7554/eLife.61666

Myers, J. C., Smith, E. H., Leszczynski, M., O’Sullivan, J., Yates, M. J., McKhann, G., Mesgarani, N., Schroeder, C., Schevon, C., & Sheth, S. A. (2022). The spatial reach of neuronal coherence and spike-field coupling across the human neocortex [Research Articles, Behavioral/Cognitive]. Journal of Neuroscience, 42(32), 6285–6294. 10.1523/JNEUROSCI.0050-22.2022

Nazmuddin, M., Philippens, I. H. C. H. M., & van Laar, T. (2021). Electrical stimulation of the nucleus basalis of Meynert: A systematic review of preclinical and clinical data. Scientific Reports, 11, 11751. 10.1038/s41598-021-91391-0

Orta-Salazar, E., Feria-Velasco, A. I., & Díaz-Cintra, S. (2019). Primary motor cortex alterations in Alzheimer disease: A study in the 3xTg-AD model. Neurología (English Edition), 34(7), 433–439. 10.1016/j.nrleng.2019.02.001

Pedreira, C., Martinez, J., Ison, M. J., & Quian Quiroga, R. (2012). How many neurons can we see with current spike sorting algorithms? Journal of Neuroscience Methods, 211(1), 58–65. 10.1016/j.jneumeth.2012.07.010

Picton, B., Wong, J., Lopez, A. M., Solomon, S. S., Andalib, S., Brown, N. J., Dutta, R. R., Paff, M. R., Hsu, F. P., & Oh, M. Y. (2024). Deep brain stimulation as an emerging therapy for cognitive decline in Alzheimer disease: Systematic review of evidence and current targets. World Neurosurgery, 184, 253–266.e2. 10.1016/j.wneu.2023.12.083

Poorthuis, R. B., Enke, L., & Letzkus, J. J. (2014). Cholinergic circuit modulation through differential recruitment of neocortical interneuron types during behaviour. The Journal of Physiology, 592(Pt 19), 4155–4164. 10.1113/jphysiol.2014.273862

Ríos, A. S., Oxenford, S., Neudorfer, C., et al. (2022). Optimal deep brain stimulation sites and networks for stimulation of the fornix in Alzheimer’s disease. Nature Communications, 13(1), 7707. 10.1038/s41467-022-34510-3

Rothlind, J. C., York, M. K., Carlson, K., et al.; CSP-468 Study Group. (2015). Neuropsychological changes following deep brain stimulation surgery for Parkinson’s disease: Comparisons of treatment at pallidal and subthalamic targets versus best medical therapy. Journal of Neurology, Neurosurgery & Psychiatry, 86(6), 622–629. 10.1136/jnnp-2014-308119

Roquet, D., Noblet, V., Anthony, P., Philippi, N., Demuynck, C., Cretin, B., Martin-Hunyadi, C., Loureiro de Sousa, P., & Blanc, F. (2017). Insular atrophy at the prodromal stage of dementia with Lewy bodies: A VBM DARTEL study. Scientific Reports, 7, Article 9437. 10.1038/s41598-017-09884-7

Saalmann, Y. B., Pigarev, I. N., & Vidyasagar, T. R. (2007). Neural mechanisms of visual attention: How top-down feedback highlights relevant locations. Science, 316(5831), 1612–1615. 10.1126/science.1139140

Suvà, D., Favre, I., Kraftsik, R., Esteban, M., Lobrinus, A., & Miklossy, J. (1999). Primary motor cortex involvement in Alzheimer disease. Journal of Neuropathology & Experimental Neurology, 58(11), 1125–1134. 10.1097/00005072-199911000-00002

Sarter, M., Parikh, V., & Howe, W. M. (2009). Phasic acetylcholine release and the volume transmission hypothesis: Time to move on. Nature Reviews Neuroscience, 10(5), 383–390. 10.1038/nm2635

Takata, N., Mishima, T., Hisatsune, C., Nagai, T., Ebisui, E., Mikoshiba, K., & Hirase, H. (2011). Astrocyte calcium signaling transforms cholinergic modulation to cortical plasticity in vivo. Journal of Neuroscience, 31(49), 18155–18165. 10.1523/JNEUROSCI.5289-11.2011

van der Zande, J. J., Gouw, A. A., van Steenoven, I., Scheltens, P., Stam, C. J., & Lemstra, A. W. (2018). EEG characteristics of dementia with Lewy bodies, Alzheimer’s disease and mixed pathology. Frontiers in Aging Neuroscience, 10, 190. 10.3389/fnagi.2018.00190

van Ede, F., Quinn, A. J., Woolrich, M. W., & Nobre, A. C. (2018). Neural oscillations: Sustained rhythms or transient burst-events? Trends in Neurosciences, 41(7), 415–417. 10.1016/j.tins.2018.04.004

Verma, A. K., Nandakumar, B., Acedillo, K., Yu, Y., Marshall, E., Schneck, D., Fiecas, M., Wang, J., MacKinnon, C. D., Howell, M. J., Vitek, J. L., & Johnson, L. A. (2024). Slow-wave sleep dysfunction in mild parkinsonism is associated with excessive beta and reduced delta oscillations in motor cortex. Frontiers in Neuroscience, 18, Article 1338624. 10.3389/fnins.2024.1338624

Vejmola, Č., Tylš, F., Piorecká, V., Koudelka, V., Kadeřábek, L., Novák, T., & Páleníček, T. (2021). Psilocin, LSD, mescaline, and DOB all induce broadband desynchronization of EEG and disconnection in rats with robust translational validity. Translational Psychiatry, 11, Article 506. 10.1038/s41398-021-01603-4

Vidyasagar, T. R., & Levichkina, E. (2019). An integrated neuronal model of claustral function in timing the synchrony between cortical areas. Frontiers in Neural Circuits, 13, 3. 10.3389/fncir.2019.00003

Wang, Y. L., Avigdor, T., Hannan, S., Abdallah, C., Dubeau, F., Peter-Derex, L., & Frauscher, B. (2024, April 17). Intracerebral dynamics of sleep arousals: A combined scalp–intracranial EEG study. Journal of Neuroscience, 44(16), e0617232024. 10.1523/JNEUROSCI.0617-23.2024

Xu, M., Chung, S., Zhang, S., Zhong, P., Ma, C., Chang, W.-C., … Dan, Y. (2015). Basal forebrain circuit for sleep-wake control. Nature Neuroscience, 18(11), 1641–1647. 10.1038/nn.4143

Xu, J., Liu, B., Shang, G., Liu, S., Feng, Z., Yang, H., Liu, D., Chang, Q., Chen, Y., Yu, X., & Mao, Z. (2024). Deep brain stimulation of the nucleus basalis of Meynert in severe Alzheimer’s disease. Journal of Alzheimer’s Disease Reports, 8(1), 1573–1586. 10.1177/25424823241296780

Wise, M. S., Arand, D. L., Auger, R. R., Brooks, S. N., & Watson, N. F. (2007). Treatment of narcolepsy and other hypersomnias of central origin: An American Academy of Sleep Medicine review. Sleep, 30(12), 1712–1727. 10.1093/sleep/30.12.1712

Yu, Y., Liu, F., Wang, W., & Lee, T. S. (2004). Optimal synchrony state for maximal information transmission. NeuroReport, 15(10), 1605–1610. 10.1097/01.wnr.0000134993.81804.22

Zhang, Y., Ren, R., Yang, L., Zhang, H., Shi, Y., Okhravi, H. R., Vitiello, M. V., Sanford, L. D., & Tang, X. (2022). Sleep in Alzheimer’s disease: A systematic review and meta-analysis of polysomnographic findings. Translational Psychiatry, 12(1), 136.

Zhang, N. K., Zhang, S. K., Zhang, L. I., Tao, H. W., & Zhang, G. W. (2023). Sensory processing deficits and related cortical pathological changes in Alzheimer’s disease. Frontiers in Aging Neuroscience, 15, 1213379. 10.3389/fnagi.2023.1213379

